# Cerebra: a computationally efficient framework for accurate protein structure prediction

**DOI:** 10.1101/2024.02.02.578551

**Authors:** Jian Hu, Weizhe Wang, Bo Zhang, Yike Cao, Haipeng Gong

**Author notes:** These authors contributed equally to this work (Jian Hu, Weizhe Wang).

## Abstract

Remarkable progress has been made in the field of protein structure prediction. Representative methods like AlphaFold and RoseTTAFold achieve prediction accuracy close to experimental structural determination, but at the cost of heavy computational consumption for model training. In this work, we propose a new framework, Cerebra, for improving the computational efficiency of protein structure prediction. In this network architecture, multiple sets of atomic coordinates are predicted parallelly and their mutual complementarity is leveraged to rapidly improve the quality of predicted structures through a new attention mechanism named Path Synthesis Attention (PSA). Consequently, Cerebra markedly reduces the computational consumption, achieving evident acceleration on both model training and inference, in comparison to OpenFold, the academic version of AlphaFold2. When evaluated on the CAMEO 2025 dataset, the Cerebra model trained on a limited number of GPUs shows a comparable performance to the published OpenFold model that has been thoroughly optimized using a plethora of computational resources. Moreover, ablation studies confirm that the PSA mechanism efficiently accelerates the network convergence, suggesting its potential in handling the many-body effects of protein geometries in graph-based models. Finally, the fast inference of Cerebra effectively assists hallucination-based *de novo* protein backbone design, allowing the production of abundant backbone structures with sufficient designability and novelty. Cerebra is freely available at https://github.com/Gonglab-THU/Cerebra, with a web server accessible at https://structpred.life.tsinghua.edu.cn/server_cerebra.html.

## Introduction

As essential components of the living systems, proteins carry out highly diversified biological functions by folding into native structures. Experimental structural determination methods are, however, time-consuming and labor-intensive, incapable of fulfilling the vast demands on protein structures. With the rapid accumulation of biological data and the unprecedented advancement in artificial intelligence, AlphaFold2^1^ has made breakthroughs in protein structure prediction, achieving a prediction accuracy close to experimental resorts on most protein monomers, and therefore has become a fundamental research tool in life sciences.

The success of AlphaFold2 has opened up new possibilities for substantial downstream explorations, such as protein complex structure prediction^2–4^, protein-ligand/small molecule interaction prediction^5–7^, protein property prediction^8–11^, and prediction of protein mutational effects on pathogenicity^12^. The model architectures of AlphaFold2, particularly the Structure Module and its subcomponents (*e*.*g*., the Invariant Point Attention (IPA) motif) that are designed for atomic coordinate modeling, have been repeatedly applied with only minor modifications in related fields like single-sequence-based protein structure prediction^13–15^ and diffusion-based protein structure generation^16,17^. In hallucination-based *de novo* protein design, the pre-trained AlphaFold2 model has been conversely used as a scoring function to constrain the sequence generators, which prominently improves the designability of produced protein backbones^18–20^. RoseTTAFold^21,22^ is the only available framework that adopts a significantly different Structure Module from AlphaFold2 but achieves a similar level of high-accuracy protein structure prediction. Similarly, this framework has demonstrated its capability in improving the prediction of protein-protein interaction, protein-nucleic acid binding, and protein-small molecule binding^23^, as well as in effectively guiding the diffusion-based protein structure generation in RFdiffusion^24^. Recently, state-of-the-art all-atom structure predictors, exemplified by AlphaFold3^25^ as well as other rival systems^26–30^, have upgraded the Structure Module in the AlphaFold2 framework by a diffusion-based architecture, thereby providing a unified framework for predicting complexes composed of proteins, nucleic acids, small molecules, ions, and modified residues. Albeit successful, the diffusion module remarkably increases the model inference time due to its iterative sampling nature, which hinders the implementation of such structure predictors as alternative substitutes of AlphaFold2 in facilitating hallucination-based *de novo* protein design.

Despite the successes of AlphaFold and RoseTTAFold and their profound impacts on protein design and engineering, both frameworks still suffer from a severe limitation, namely, the enormous consumption of GPU resources required for model training. For instance, OpenFold^31^, the academic implementation of AlphaFold2, requires approximately 50,000 GPU hours to train a high-accuracy model. Such a heavy computational burden restricts the continual updating of model parameters with newly available data and hinders independent research groups from extensively investigating model interpretability, architectural optimization, and integration with downstream tasks. This problem is further exacerbated in diffusion-based generative structure predictors like AlphaFold3, where iterative sampling and stochastic generation not only impose additional computational costs but may lead to unwanted variability in the predicted structures of challenging targets. Therefore, designing a computationally more efficient framework than AlphaFold and RoseTTAFold, particularly a non-diffusion-based Structure Module capable of performing parallel coordinate prediction, remains an important direction for further exploration.

In this work, to address this issue, we developed a protein structure prediction framework named Cerebra (<u>c</u>o<u>e</u>volution of <u>r</u>esidue <u>e</u>mbedding and <u>b</u>etween-<u>r</u>esidue <u>a</u>ttention) by designing an innovative Structure Module to enhance the computational efficiency. The Cerebra model has two major characteristics. Firstly, it predicts multiple sets of atomic coordinates simultaneously to reach a parallelized training effect, speeding up model convergence and reducing training costs significantly. Specifically, when trained on the same dataset containing 10,000 proteins, Cerebra attains a training acceleration of sixfold over OpenFold. Secondly, the Structure Module allows accurate prediction of various local portions of the target protein within the multiple sets of atomic coordinates and the complementarity between these local structural motifs is then leveraged by a novel attention mechanism called Path Synthesis Attention (PSA) to further suppress prediction errors during the reconstruction of the global structure, a strategy that effectively reduces the number of folding steps for protein structure modeling and improves the inference efficiency of protein structure prediction. Through model training on limited computational resources, Cerebra achieves an average TM-score of 0.860 on the 573 CAMEO targets collected over the year of 2025, reaching a comparable performance to OpenFold, with only slight inferiority to RoseTTAFold2 and AlphaFold3. Furthermore, we delved into the PSA module mechanistically and validated its effectiveness in accelerating the network convergence and in improving the prediction of residue-frame translations and rotations. Finally, we repurposed Cerebra for hallucination-based *de novo* protein backbone design. Benefiting from Cerebra’s high inference efficiency, our approach successfully generates novel protein backbones of high modeling confidence and designability across a range of sequence lengths under affordable computational consumption.

## Methods

### Overview of the Cerebra framework

As shown in Figure 1a, Cerebra consists of three components: an Embedding Module for processing raw data, an Evoformer Module for integrating 1D and 2D features, and a Structure Module for generating protein structures. The basic architecture of the Embedding Module is similar to that of AlphaFold2, which integrates the multiple sequence alignment (MSA) with the recycling feedback from the preceding cycle. Here, we exclude the extra MSA and template inputs in AlphaFold2 but instead employ the ESM-2^13^ embedding of the target sequence as a controllable additional input, selectively strengthening this signal for targets with sparse MSAs. The Embedding Module generates sequence and pairwise features, which are referred to as 1D and 2D features, respectively, in our model. The Evoformer Module is directly implanted from AlphaFold2 for the mutual update of 1D and 2D features, but with a simplification from the original 48 copies of Evoformer motifs to 32 copies to fit in the limited GPU memory (Figure S2).

**Figure 1.**
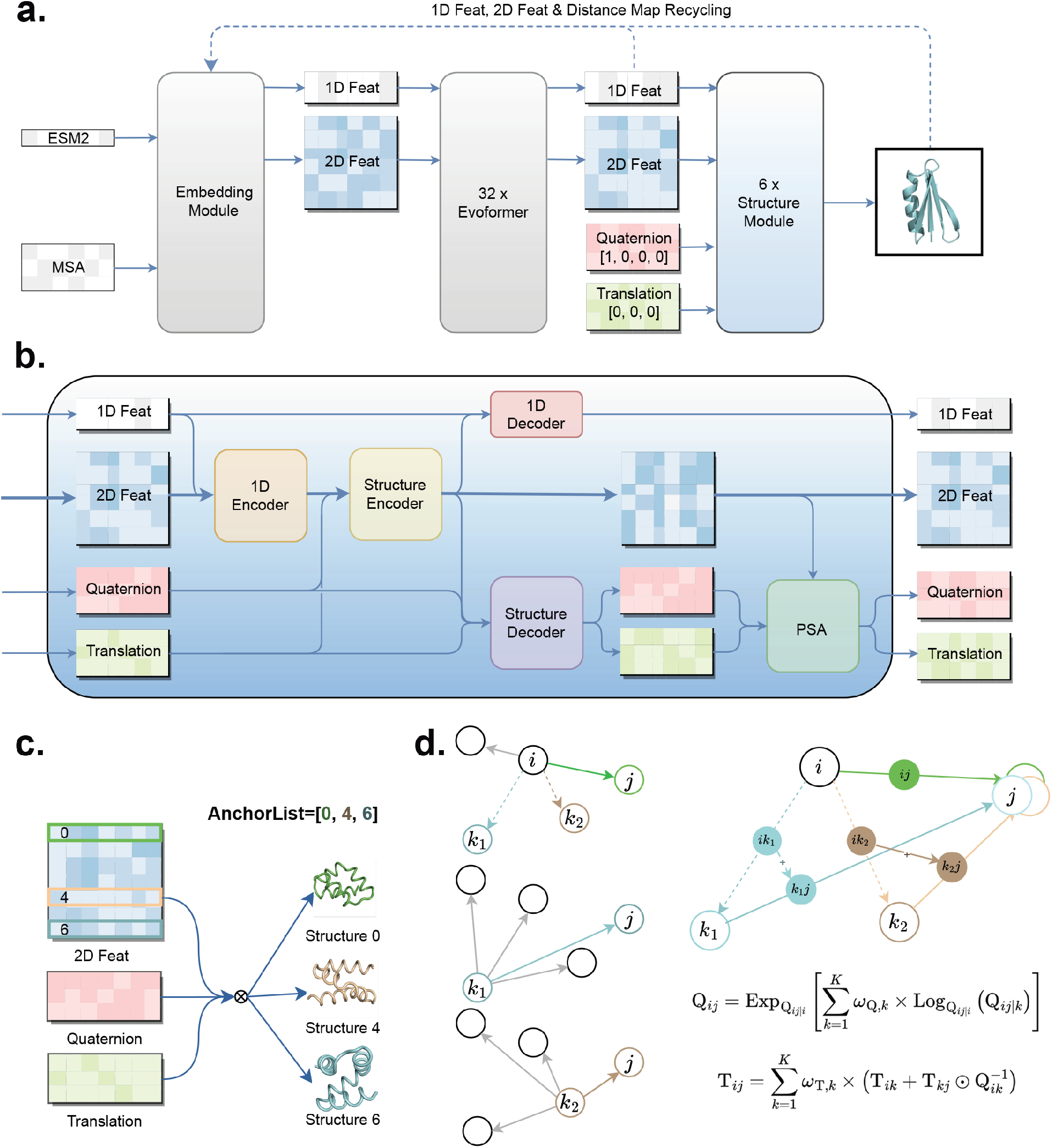
The schematic representation of the architecture of the Cerebra model. **a)** The model architecture of Cerebra. **b)** The expanded architecture of one structure generation motif in the Structure Module. **c)** Based on the AnchorList, for each anchor residue, its corresponding row of the 2D features is used to update the translations and rotations of all residues and thus construct the structure within this reference frame. **d)** The PSA module adjusts the translation and rotation of residue *j* in the local coordinate system of residue *i* by integrating coordinate information from all paths connecting *i* and *j* through no more than one intermediate anchor residues (*k*_1_, *k*_2_, etc.), a strategy equivalent to recombining the multiple predicted structures for structural refinement.

In this work, we designed a novel Structure Module (Figure 1b) by sequentially stacking 6 structure generation motifs without weight sharing, each of which takes as input the 1D features, 2D features and *K* sets of translations (in vectors) and rotations (in quaternions), where the hyper-parameter *K* stands for the number of anchor residues whose local coordinate systems are used as reference frames for structural generation and evaluation. Here, the local coordinate system of each amino acid residue is constructed following the AlphaFold2 definition^1^. After a series of operations, the updated 1D features, 2D features and *K* sets of translations/rotations are then fed into the next structure generation motif. Notably, we use “translation” (symbolized as T) and “quaternion” (symbolized as Q) to refer to the origins and rotation matrices of the local coordinate systems of all residues within each of the *K* reference frames. The 1^*st*^ structure generation motif reads in the 1D and 2D features from the Evoformer Module and initializes the translation and quaternion values as [0, 0, 0] and [1, 0, 0, 0], respectively.

Unlike AlphaFold2 and RoseTTAFold2 that rely on the 1D embedding for structure generation, Cerebra updates the *K* sets of translations and quaternions based on the 2D features within each structure generation motif. Specifically, for each of the *K* anchor residues, the structure generation motif employs its corresponding row within the 2D features to predict the changes of translations and rotations for all residues in this reference frame. To reduce computational complexity, the *K* anchor residues are distributed roughly uniformly along the sequence, composing the anchor residue list (Figure 1c). Structures corresponding to the *K* sets of translations and quaternions are then recombined by the PSA module (Figure 1d) to further reduce prediction errors.

### Encoders and decoders in the Structure Module

In Cerebra, the 2D features play a central role in the Structure Module. Specifically, in each structure generation motif, all features are first fused with the original 2D features in the encoding process and the updated 2D features are then distributed to the 1D features and structure features (*i*.*e*. translations and quaternions) in the decoding process (Figure 1b). The encoding process is accomplished by the 1D encoder (Algorithm 7) and the structure encoder (Algorithm 8). The former produces attention maps through self-attention on the 1D features and then integrates with the original 2D features for information updating, whereas the latter collects the C_*α*_ and C_*β*_ inter-residue distance distributions as well as the normalized translations and quaternions to update the 2D features. Similarly, the decoding process is accomplished by the 1D decoder (Algorithm 9) and the structure decoder (Algorithm 10). The former transforms the updated 2D features into attention maps, from which the 1D features are updated through the attention mechanism, whereas the latter predicts the changes of translations and quaternions for all residues in the reference frame of each anchor residue using the corresponding row of the updated 2D features (Figure 1c).

### Path Synthesis Attention (PSA) module in the Structure Module

We obtain the predicted translations and rotations of all residues in the reference frames of anchor residues (*i, k*_1_, *k*_2_, etc. in Figure 1d) from the structure decoder. To infer the translation of residue *j* in the local coordinate system of residue *i*, we can either use the raw prediction value T_*ij*_ directly, or integrate the information from reference frames 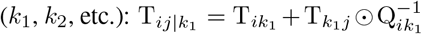 and 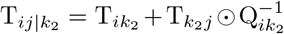, where ⊙ is a quaternion operator, whose definition can be found in Supplementary Information 3.2 (see Algorithm 3 for the implementation). The rotation (*i*.*e*. quaternion) of residue *j* could be derived directly from Q_*ij*_ or using similar formulas: 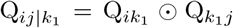 and 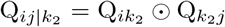 . More generally, we can refer to a group of three residues, *i*.*e*. the source anchor residue *i*, an intermediate anchor residue *k* and the target residue *j*, to compose a path (*i* → *k* → *j*) for the estimation of translation and rotation of residue *j* in the reference frame of residue *i*, as in Equations 1 and 2:

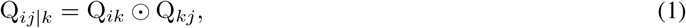

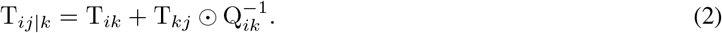

Since our network engages *K* anchor residues, there exist a total of *K* paths for updating the translation and rotation of residue *j* relative to residue *i*, which also include the raw prediction when the intermediate residue *k* is set as *i*. Considering the inevitable errors in regression prediction by the neural networks, we take a weighted average of values obtained from all paths connecting residues *i* and *j* to obtain a more accurate prediction for the translation and rotation of residue *j* in the reference frame of residue *i*. The weighted averaging is accomplished by the PSA module through a novel attention mechanism (Algorithm 12). Notably, the PSA module is employed independently for translations and rotations, with different learnable parameters. Hence, we denote the corresponding paths as translation paths and rotation paths, respectively.

We first apply interpolation in the Lie algebra of rotations to realize rotation updating. Specifically, the weights *ω*_Q,*k*_ are derived from the 2D features and quaternions (see Figure S16 for a schematic description). To avoid directly averaging quaternions on the unit sphere, we use the direct rotation path Q_*ij*|*i*_ as the base quaternion for residue *j* in the reference frame of residue *i*, where Q_*ij*|*i*_ = Q_*ii*_ ⊙ Q_*ij*_ = Q_*ij*_. Each path quaternion is sign-aligned to the base quaternion as 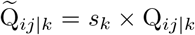 where *s*_*k*_ = 1 if the inner product ⟨Q_*ij*|*k*_, Q_*ij*|*i*_⟩ ≥ 0 and *s*_*k*_ = −1 otherwise. The aligned relative rotations are then mapped to the tangent space by the logarithmic map, averaged with the attention weights, and mapped back by the exponential map, as in Equation 3:

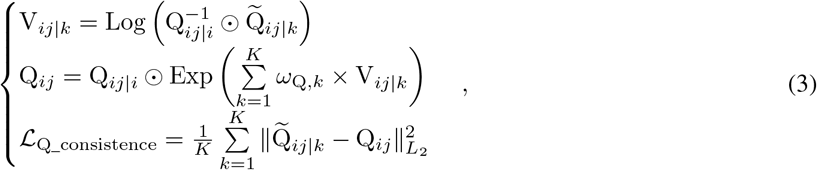

where Log and Exp denote the logarithmic and exponential maps between the quaternion representation of rotations and its corresponding Lie algebra representation (Algorithm 4; see Supplementary Information 3.3 for details). We also calculate the squared *L*_2_ distances between the sign-aligned quaternions estimated from different paths and the final value, and take the average to compose an additional consistency loss *L*_Q_consistence_ to enforce small variation among rotations updated along different paths.

The updated rotations can be applied to the translation calculations subsequently through coupled translational and rotational transformations. Similar to the weight calculation method for rotation paths, we integrate all translation paths together, but without back-and-forth transformations between Lie group and Lie algebra, as in Equation 4:

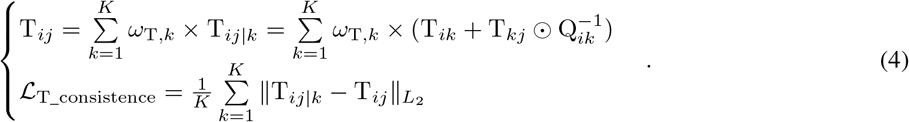

For translations, we use the *L*_2_ norm in the consistency loss L_T_consistence_ to constrain the variation among translations updated along different paths.

### Datasets and metrics for model training and evaluation

#### Training dataset

We used the Protein Data Bank (PDB) dataset and the AlphaFold2 distillation dataset constructed by the OpenFold consortium as the training data. The original OpenFold training set contains approximately 130,000 protein monomers from the PDB and 270,000 protein monomers from the AlphaFold Protein Structure Database (AFDB). In addition, the newly released OpenFold3^32^ provides AlphaFold2-predicted structures for approximately 60,000 intrinsically disordered proteins, which were also incorporated into our training set. After excluding samples of length < 50 residues, we retained ∼116,000 PDB protein samples for model training and further incorporated ∼268,000 proteins from the AlphaFold2 distillation dataset together with ∼46,000 disordered proteins with AlphaFold2-predicted structures for fine-tuning at the late training stage. The input features include the MSA raw data and the ESM-2 embedding of the target sequence, without structural templates (see Supplementary Information 6 for details).

#### Test dataset

Since the OpenFold dataset contains PDB files released by December 2021, we used the annual CAMEO dataset from the year of 2025 as the primary test set to validate the effectiveness of our model. After data cleaning and excluding samples of length < 50 residues, we retained 573 protein samples in the CAMEO test set. The MSAs for all evaluation targets were generated using the MSA-search pipeline implemented in LocalColabFold^33^, and the resulting MSA files were used identically as inputs to Cerebra and OpenFold.

#### Evaluation metrics

We engaged TM-score^34^ and lDDT^35^ to evaluate the quality of predicted structures, in terms of the global topology and local details, respectively.

### Hallucination-based *de novo* protein backbone design

We repurposed Cerebra for *de novo* protein backbone design by initializing each optimization trajectory from a random amino acid sequence. Cerebra predicted the protein backbone and residue-level pLDDT values, and the sequence *s* of length *L* was optimized by minimizing the complement of the mean modeling confidence,

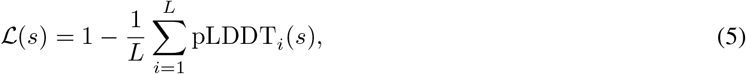

where pLDDT values were normalized to the range [0, 1]. Sequence mutations were accepted using Markov Chain Monte Carlo (MCMC) sampling with simulated annealing. We developed two protocols: the refined protocol employs a gradual 5,000-step search, whereas the fast protocol preferentially mutates low-confidence regions and thus reduces the optimization to 1,500 steps. In the final evaluation, we chose the refined protocol for proteins of 50 and 100 residues, but adopted the fast protocol for proteins of 150 and 200 residues. No structural template or target topology was provided during optimization.

For each generated backbone, ProteinMPNN^36^ was used to design amino acid sequences, which were subsequently refolded with ESMFold^13^. Backbone root-mean-square-distance (RMSD) and TM-score between the refolded and generated structures were used to evaluate designability. Structural novelty was assessed by searching the generated backbones against the PDB with Foldseek^37^.

Detailed optimization protocols and evaluation settings are provided in Supplementary Information 8.

## Results

### Model performance evaluation

Cerebra was trained using the publicly available training datasets released by the OpenFold consortium. Since Open-Fold is an open-source implementation of AlphaFold2 with comparable state-of-the-art performance, we selected the officially released OpenFold model as the primary benchmark for the fair evaluation of Cerebra. We additionally compared Cerebra with RoseTTAFold2 and AlphaFold3, two state-of-the-art protein structure prediction models that utilize a large scale of additional macromolecules for model training. All models were evaluated on the cleaned 2025 CAMEO test set comprising 573 targets, and prediction accuracy was assessed using TM-score and RMSD.

As shown in Figure 2a&b, Cerebra achieves a comparable performance to OpenFold, with nearly identical mean TM-scores (0.860 for Cerebra vs. 0.861 for OpenFold) and mean RMSD values (4.042 Å for Cerebra vs. 4.055 Å for OpenFold). The per-target comparison further reveals a broad agreement between Cerebra and OpenFold, while also identifying targets for which Cerebra produces substantially more accurate predictions (Figure 2c). Overall, Cerebra outperforms OpenFold (in TM-score) on 221 targets, among which the lead is greater than 0.1 for 27 targets. Figure 2d presents two exemplar targets, 9BGY and 9FW7, where Cerebra attains TM-scores of 0.947 and 0.954, respectively, in comparison with 0.615 and 0.344 by OpenFold. These results demonstrate that Cerebra reaches an accuracy comparable to the officially released OpenFold model and can provide complementary high-quality predictions for individual targets. Although RoseTTAFold2 and AlphaFold3 exhibit slightly higher performance than Cerebra and OpenFold (in both the mean TM-scores and mean RMSD), the superiority of the former is likely attributable to the recruitment of a large scale of AFDB structures (tens of millions) for model training, while the advantage of the latter possibly arises from the advanced diffusion-module-based design, which effectively utilizes molecules other than proteins in the PDB to improve training at the sacrifice of inference speed.

**Figure 2.**
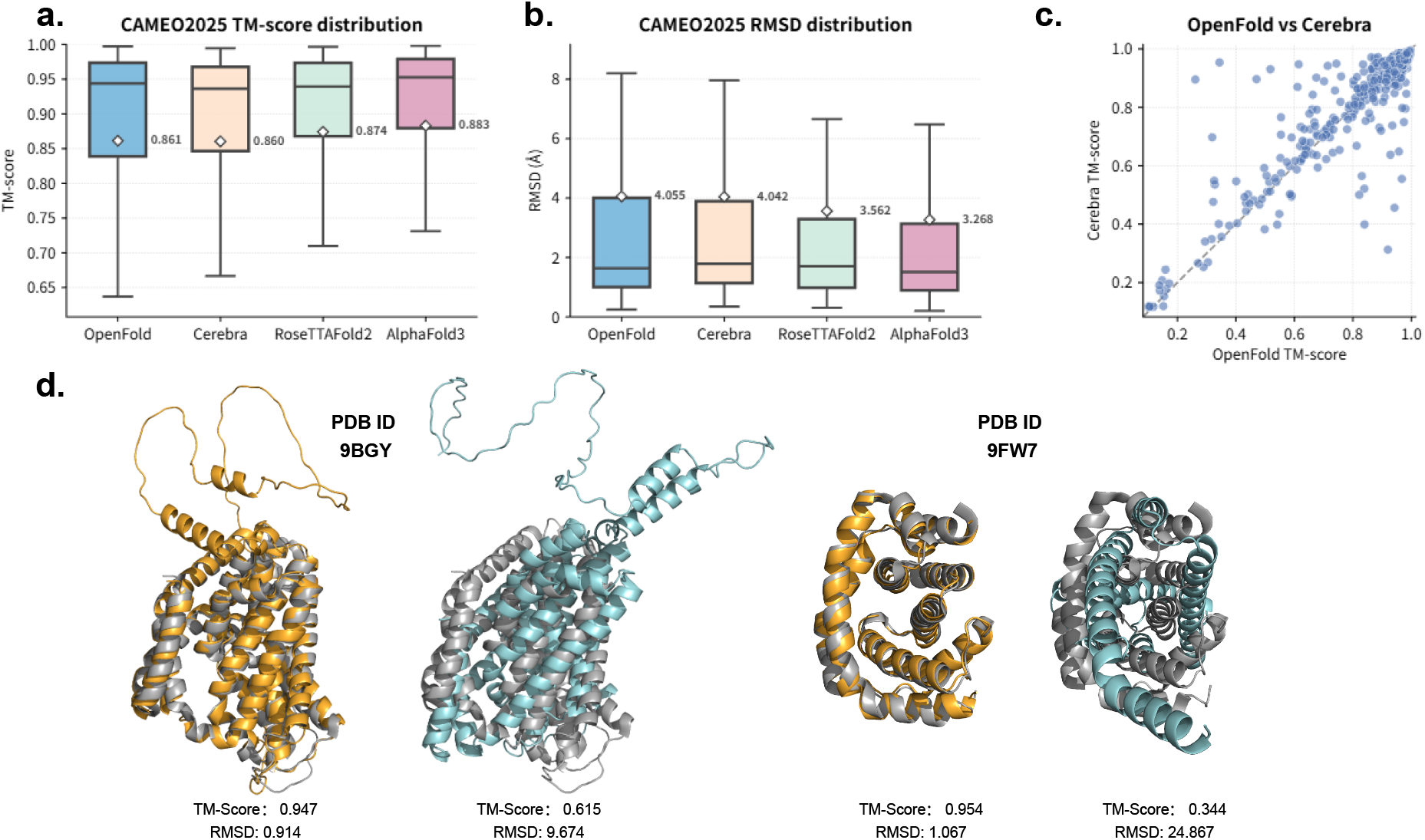
Comparison of protein structure prediction performance on the CAMEO 2025 test set. **a–b)** Distributions of TM-score (**a**) and RMSD (**b**) for structures predicted by OpenFold, Cerebra, RoseTTAFold2, and AlphaFold3 on 573 CAMEO targets. Center lines represent medians, boxes indicate interquartile ranges, and whiskers extend to the most extreme values within 1.5 times the interquartile range. White diamonds and adjacent values indicate means. **c)** Per-target comparison of TM-scores obtained by Cerebra and OpenFold. The dashed line indicates equal performance. **d)** Two representative targets 9BGY and 9FW7, for which Cerebra outperforms OpenFold. Cerebra and OpenFold predictions are colored orange and cyan, respectively, and are structurally aligned to the experimentally derived structures colored in gray.

Besides the above evaluation, we also benchmarked our Cerebra model on the CAMEO 2024 dataset, which contains 643 protein targets collected in the preceding year, and reached a similar conclusion. That is, Cerebra achieves a comparable performance to OpenFold, with complementarity in individual targets. The corresponding results are summarized in Supplementary Information 1.1.

### Analysis on the training and inference efficiency

One of the prominent advantages of Cerebra is its ability to approach high prediction accuracy through model training using significantly less computational resources. Here, we benchmarked the difference in training costs between Cerebra and OpenFold by deriving their training curves (*i*.*e*. increment of model performance with training time or steps) on a small dataset of ∼10,000 protein chains (named as the 10K dataset, see Supplementary Information 6 for details). For a fair comparison with OpenFold, we blocked the ESM-2 embedding and retained the MSA as the only input feature for Cerebra. As shown in Figure S6a, in such a scenario, the standard Cerebra model (*i*.*e*. the Cerebra model with 32 anchor residues) attains training lDDT scores of 0.6 and 0.7 at the time points of 21 and 34 GPU hours, respectively, whereas OpenFold takes 91 and 205 GPU hours to reach the same lDDT scores, respectively. Hence, Cerebra exhibits a speed-up of model training by about 6 folds when using the same input features and labels extracted from the completely identical training set. Adding pre-trained ESM-2 embeddings to Cerebra further reduces the corresponding training time to 14 and 24 GPU hours, respectively, representing an approximately 8.5-fold acceleration relative to OpenFold. As for the training steps, Cerebra requires half of the update steps of OpenFold to achieve the same training effect (Figure S6b).

We next evaluated the inference efficiency of Cerebra, OpenFold and AlphaFold3 for proteins ranging from 100 to 1,000 residues at intervals of 100 residues. All three models were evaluated locally on a single NVIDIA A100 GPU with 80 GB of memory, with each measurement repeated three times. The model-specific inference parameters were retained at their respective official default settings. We measured the end-to-end inference time, *i*.*e*. excluding the time required for MSA search and model precompilation. Cerebra was consistently faster than both OpenFold and AlphaFold3 at all tested sequence lengths (Figure 3a). After averaging across all evaluated lengths, the mean inference time of Cerebra, OpenFold and AlphaFold3 is 17.42, 43.93 and 34.220 s, respectively, which correspond to an overall speedup of 2.52-fold and 1.96-fold by Cerebra relative to OpenFold and AlphaFold3, respectively. Particularly, for large targets of 1,000 residues, Cerebra completes inference in 46.41 s, compared with 123.66 s for OpenFold and 65.07 s for AlphaFold3, which correspond to a Cerebra speedup of 2.66-fold and 1.40-fold, respectively (Figure 3b).

**Figure 3.**
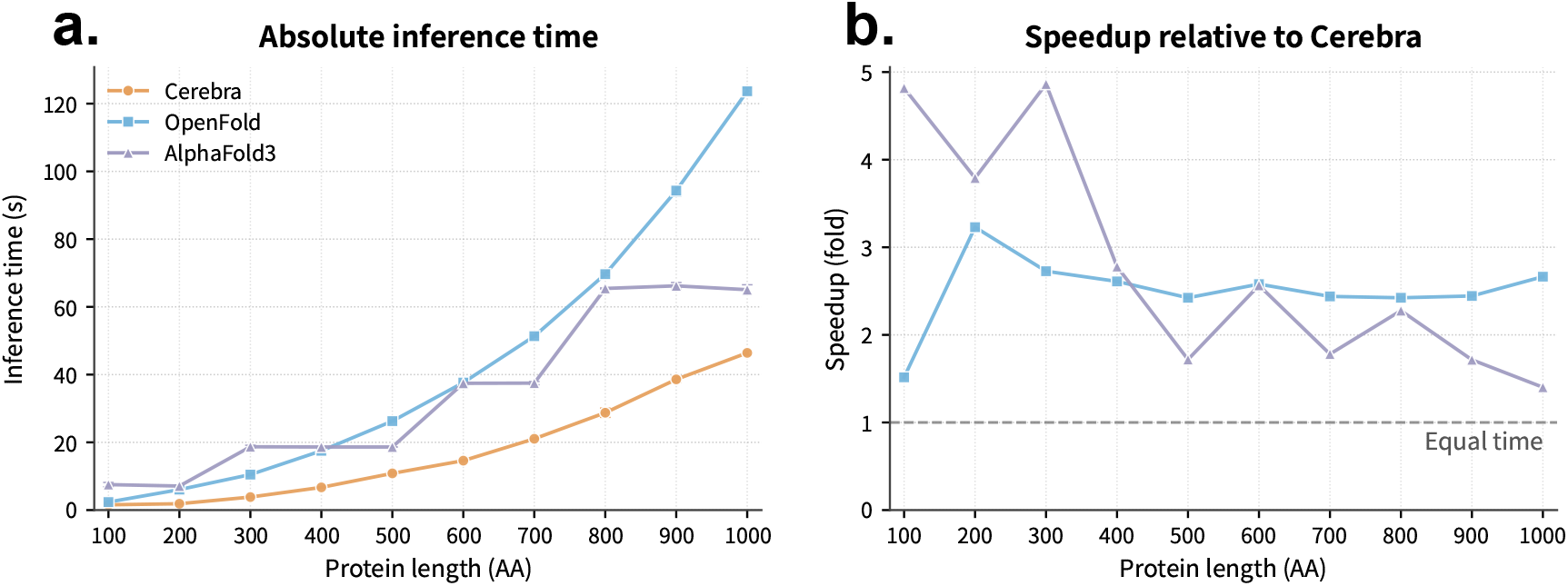
Evaluation on model inference. **a)** Comparison of the inference time (vertical axis) for Cerebra (orange), OpenFold (cyan) and AlphaFold3 (purple) on proteins of various lengths (horizontal axis). For each fixed length, the inference time is obtained from three repeats (*n* = 3). **b)** Fold of speedup by Cerebra when compared with OpenFold (cyan) and AlphaFold3 (purple), respectively. The horizontal dashed line represents the reference of no speedup.

Collectively, our experiments demonstrate that Cerebra reduces both the computational cost of model training and the time required for structure inference. Its efficient training enables ordinary research groups with limited GPU resources to independently develop, update, and fine-tune accurate structure prediction models. Meanwhile, its consistently faster inference is particularly advantageous for applications requiring intensively repeated model evaluations, including the hallucination-based *de novo* protein backbone design explored below.

### PSA accelerates training convergence

The PSA module is designed purposedly to enhance the prediction accuracy by effectively integrating the correctly predicted structural information among the multiple sets of parallelly generated atomic coordinates. Here, we conducted ablation experiments on the 10K dataset to verify the effectiveness of this module. Based on the training curves in Figure 4a, it is evident that removal of the PSA module elicits faster training at the early stage but eventually impedes the increment of model performance, with the training lDDT converging around 0.6, in sharp contrast to the standard Cerebra training process that finally approaches the lDDT score of 0.75. Similarly, if the consistency losses (*L*_Q_consistence_ in Equation 3 and *L*_T_consistence_ in Equation 4 for constraining the geometric inconsistency among generated structures) are excluded from the standard Cerebra training, the overall training process is retarded, with the training lDDT reaching a plateau of around 0.6, close to the situation of removing PSA. Furthermore, after reapplying the consistency losses to the model checkpoint obtained at approximately 14,000 steps in the above training process, the training lDDT declines first but then rises, surpassing the original checkpoint. As shown in Figure 4b, with the gradual optimization of the translational consistency loss *L*_T_consistence_, the model regains its learning capability, mimicking the standard Cerebra training process. These observations uniformly support the essential roles of the PSA module and the consistency loss functions in the optimization of Cerebra model.

**Figure 4.**
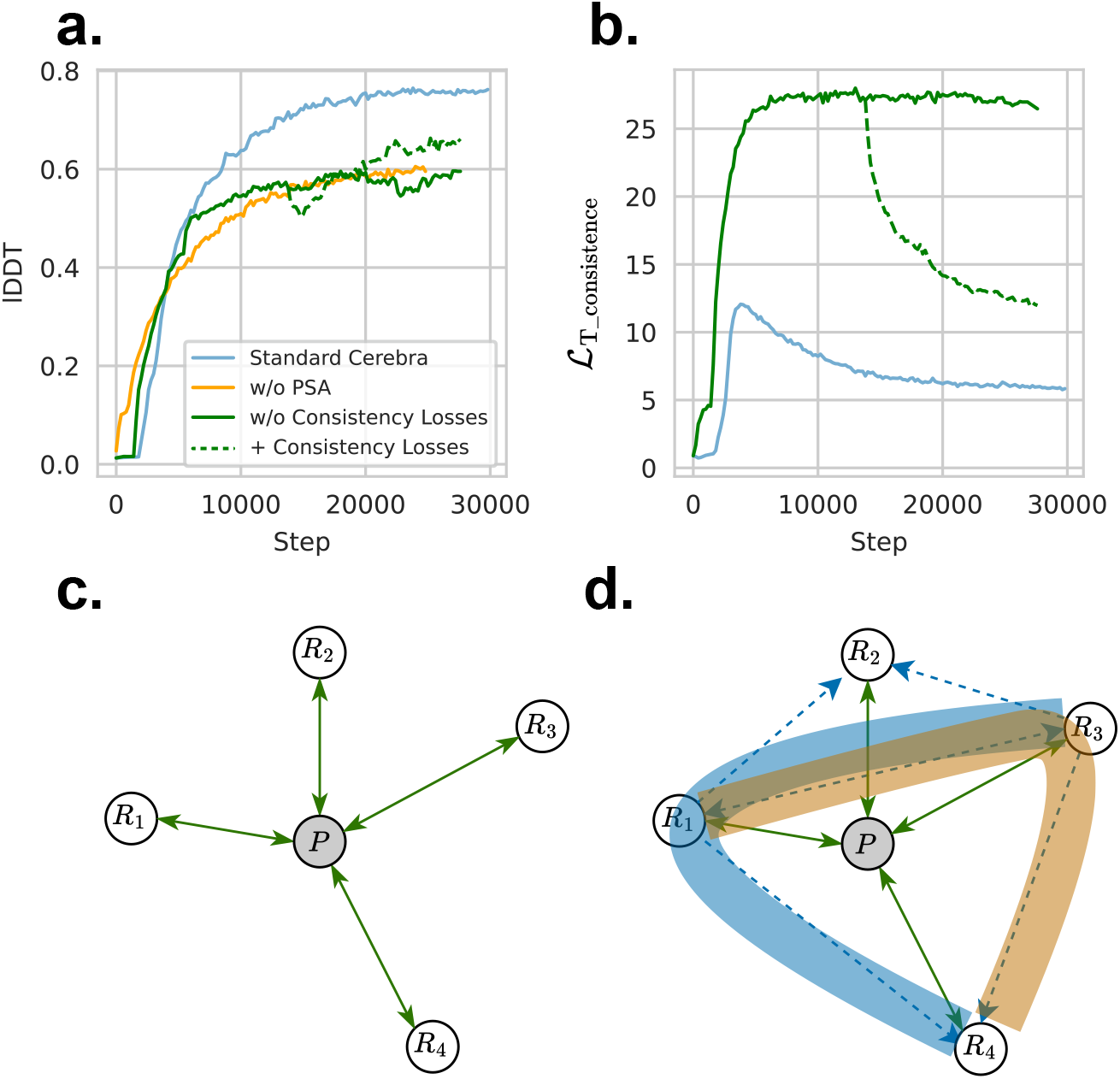
Validation on the PSA module. **a)** Training curves for the standard Cerebra model (blue line), the model with the PSA module removed (orange line), the model with the consistency loss excluded (green line), and the model with the consistency loss restored at around 14,000 steps (green dashed line). **b)** Evolution of the consistency loss for translation. **c–d)** The exchange of information between the structures generated for the reference frames of different anchor residues (*R*_1_, *R*_2_, etc.) and the hypothetical parameter space *P* . Schematic explanations are provided from a graphical viewpoint for the cases in the absence (**c**) and presence (**d**) of the PSA module.

The role of the PSA module could be more easily understood by a schematic interpretation, where each anchor residue is represented as a node (*R*_1_, *R*_2_, etc.) while the whole parameter space is simplified as a virtual node *P* . In the absence of PSA, the model training process is analogous to a graph that only allows information exchange between the residue nodes and the virtual node (Figure 4c), since parameters are independently optimized based on individual structures predicted for the reference frames of individual anchor residues. As a contrast, the PSA module enables additional information exchange among the residue nodes (Figure 4d). Suppose that the reference coordinate system is established upon the anchor residue *R*_1_ and that we hope to estimate the coordinate of residue *R*_4_, then the coordinate information of residue *R*_3_ will be integrated into that of residue *R*_4_ via the “synthesized” path 1 → 3 → 4 (brown curve) following Equation 3 and Equation 4. Similarly, exemplar “synthesized” paths 3 → 1 → 4 (blue curve) and 3 → 1 → 2 (not shown) could merge coordinate information of the intermediate residue *R*_1_ into those of other residues. Therefore, the PSA module accelerates the convergence of model training by eliciting faster and more convenient information flow across the local coordinate systems of different anchor residues. More topics on the anchor residue effects could be found in Supplementary Information 1.3.

### PSA improves the prediction on translation

The PSA module enables information integration among the multiple “synthesized” paths that are mediated by single-hop relay residues to improve the prediction of translation and rotation of residue *j* in the local coordinate system of residue *i*. To verify the impact of PSA in the translation prediction, we gathered the translation predictions T_*ij*|*k*_ and the attention weights *ω*_T,*k*_ along various translation paths (see Equation 4) generated by each structure generation motif of the Structure Module for every residue pair of *i* and *j* on the CAMEO targets, and then calculated the mean prediction errors of 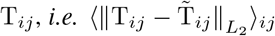 where 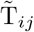 denotes the ground truth value. Here, T_*ij*_ are derived by three different approaches: 1) by the PSA weighted averaging (see Equation 4); 2) by the simple arithmetic averaging, 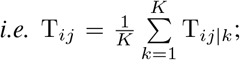 and 3) by the raw prediction, *i*.*e*. T_*ij*|*k*=*I*_. As shown in Table 1, PSA processing attains the smallest prediction error for residue pairs located at nearly all distance ranges and among nearly all Structure Module motifs. This observation demonstrates the capacity of the PSA module in improving the translation prediction by effectively integrating information from various translation paths.

**Table 1.** Prediction errors in translation by different methods.

| Residue Distance $r_{ij}$ | Method | Structure Module | | | | | |
| --- | --- | --- | --- | --- | --- | --- | --- |
|  |  | 1 | 2 | 3 | 4 | 5 | 6 |
| [0, 10) | Raw | 1.332 | 1.050 | 1.011 | 1.037 | 1.027 | 1.024 |
|  | Avg | 0.814 | **0.720** | 0.715 | 0.702 | 0.709 | 0.693 |
|  | PSA | **0.727** | 0.748 | **0.668** | **0.653** | **0.668** | **0.663** |
| [10, 20) | Raw | 2.021 | 1.637 | 1.581 | 1.595 | 1.573 | 1.567 |
|  | Avg | 1.462 | 1.237 | 1.250 | 1.218 | 1.215 | 1.227 |
|  | PSA | **1.204** | **1.171** | **1.133** | **1.113** | **1.116** | **1.127** |
| [20, 30) | Raw | 2.629 | 2.254 | 2.178 | 2.143 | 2.109 | 2.047 |
|  | Avg | 2.222 | 1.779 | 1.793 | 1.708 | 1.690 | 1.656 |
|  | PSA | **1.738** | **1.670** | **1.641** | **1.599** | **1.591** | **1.566** |
| [30, $+\infty$) | Raw | 4.735 | 4.282 | 4.145 | 3.967 | 3.910 | 3.629 |
|  | Avg | 4.665 | 3.591 | 3.573 | 3.362 | 3.300 | 3.114 |
|  | PSA | **3.537** | **3.382** | **3.331** | **3.225** | **3.195** | **3.033** |
| [0, $+\infty$) | Raw | 3.309 | 2.898 | 2.802 | 2.722 | 2.686 | 2.557 |
|  | Avg | 3.023 | 2.365 | 2.358 | 2.241 | 2.212 | 2.136 |
|  | PSA | **2.352** | **2.255** | **2.200** | **2.136** | **2.125** | **2.057** |

For each type of classification, the method showing the lowest prediction error is highlighted in bold. Raw denotes the raw prediction results, Avg refers to the simple unweighted averaging, while PSA stands for the attention-based weighted average as designed in Cerebra. $r_{ij}$ is the distance between residue $i$ and residue $j$ in the native protein structure. Prediction errors are evaluated in the unit of Å.

Next, we calculated the Pearson correlation coefficient (PCC) between the attention weights optimized by the PSA module along all translation path *ω*_T,*k*_ and the lengths of these paths in the Euclidean space (*i*.*e. l*_*ij*|*k*_ = *r*_*ik*_ + *r*_*kj*_, where *r* denotes the Euclidean distance between a pair of residues in the native structure). As shown in Table 2, the universal negative PCC between the attention weights and the path length indicates that short paths mediated by relay residue *k* sitting between the residue pair *i* and *j* are generally favored by PSA. Similarly, we evaluated the cosine values of the scalar angle of the path 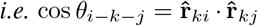 where 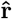 is the unit vector) and also found a universal negative PCC between this metric and the attention weights, which further corroborates the above conclusion.

**Table 2.** Pearson correlation between translation path metrics and PSA attention weights Path Metric Residue Distance *r*_*ij*_ Structure Module.

| Path Metric | Residue Distance $r_{ij}$ | Structure Module | | | | | |
| --- | --- | --- | --- | --- | --- | --- | --- |
|  |  | 1 | 2 | 3 | 4 | 5 | 6 |
| Length | [0, 10) | -0.176 | -0.260 | -0.296 | -0.282 | -0.326 | -0.255 |
|  | [10, 20) | -0.235 | -0.288 | -0.302 | -0.323 | -0.277 | -0.268 |
|  | [20, 30) | -0.212 | -0.263 | -0.275 | -0.314 | -0.244 | -0.252 |
| | [30, $+\infty$) | -0.130 | -0.174 | -0.202 | -0.217 | -0.221 | -0.190 |
| | [0, $+\infty$) | -0.174 | -0.229 | -0.253 | -0.259 | -0.250 | -0.221 |
| Angle | [0, 10) | -0.077 | -0.211 | -0.440 | -0.127 | -0.796 | -0.425 |
|  | [10, 20) | -0.216 | -0.292 | -0.469 | -0.220 | -0.718 | -0.410 |
|  | [20, 30) | -0.215 | -0.284 | -0.378 | -0.293 | -0.511 | -0.339 |
| | [30, $+\infty$) | -0.139 | -0.171 | -0.240 | -0.225 | -0.329 | -0.222 |
| | [0, $+\infty$) | -0.163 | -0.221 | -0.339 | -0.216 | -0.495 | -0.297 |

Length refers to the absolute path length in the Euclidean space, while Angle means the cosine value of the scalar angle of the path. $r_{ij}$ is the distance between residue $i$ and residue $j$ in the native protein structure. Direct paths between $i$ and $j$ (*i.e.* when $k = i$ ) are excluded from the evaluation of the Angle metric to avoid the lack of angle definition.

Here, we take 8EE0_A as an example to showcase the effects of PSA processing (Figure 5). As shown in Figure 5a, the multiple sets of atomic coordinates predicted by the 1^*st*^ Structure Module motif are highly dissimilar but become remarkably more homogeneous in structures after the PSA processing. The same effect could be observed in all downstream structure generation motifs (see the solid arrows in each column), and the quality of predicted structures is continuously enhanced with the progress through these sequentially arranged motifs (see the dashed arrows across columns). Similarly, as shown in Figure 5b, the originally broadly distributed RMSD values of the multiple sets of predicted structures relative to the ground truth (blue curves) are compressed by the PSA module, showing a sharp distribution (yellow curves) in each structure generation motif. Notably, in the last two motifs, the distributions before and after PSA processing present overlapped peaks, indicating that the sufficient improvement in the 2D features allows accurate structural modeling directly. Figure 5c shows exemplar structures predicted in the 1^*st*^ Structure Module motif. Since atomic coordinates are predicted in the reference frames of the anchor residues (see the orange spheres), coordinates of local residues close to the reference origin are well predicted while the distal portions have poor structure quality (see the red colored parts in the upper row), a typical challenging problem in regression prediction tasks using neural networks. The frustration is, however, greatly alleviated after PSA processing (see the lower row), because coordinates of the distantly positioned residues could be recovered through relay residues sitting between the target and source, by utilizing more reliable predictions on translations and rotations over shorter ranges.

**Figure 5.**
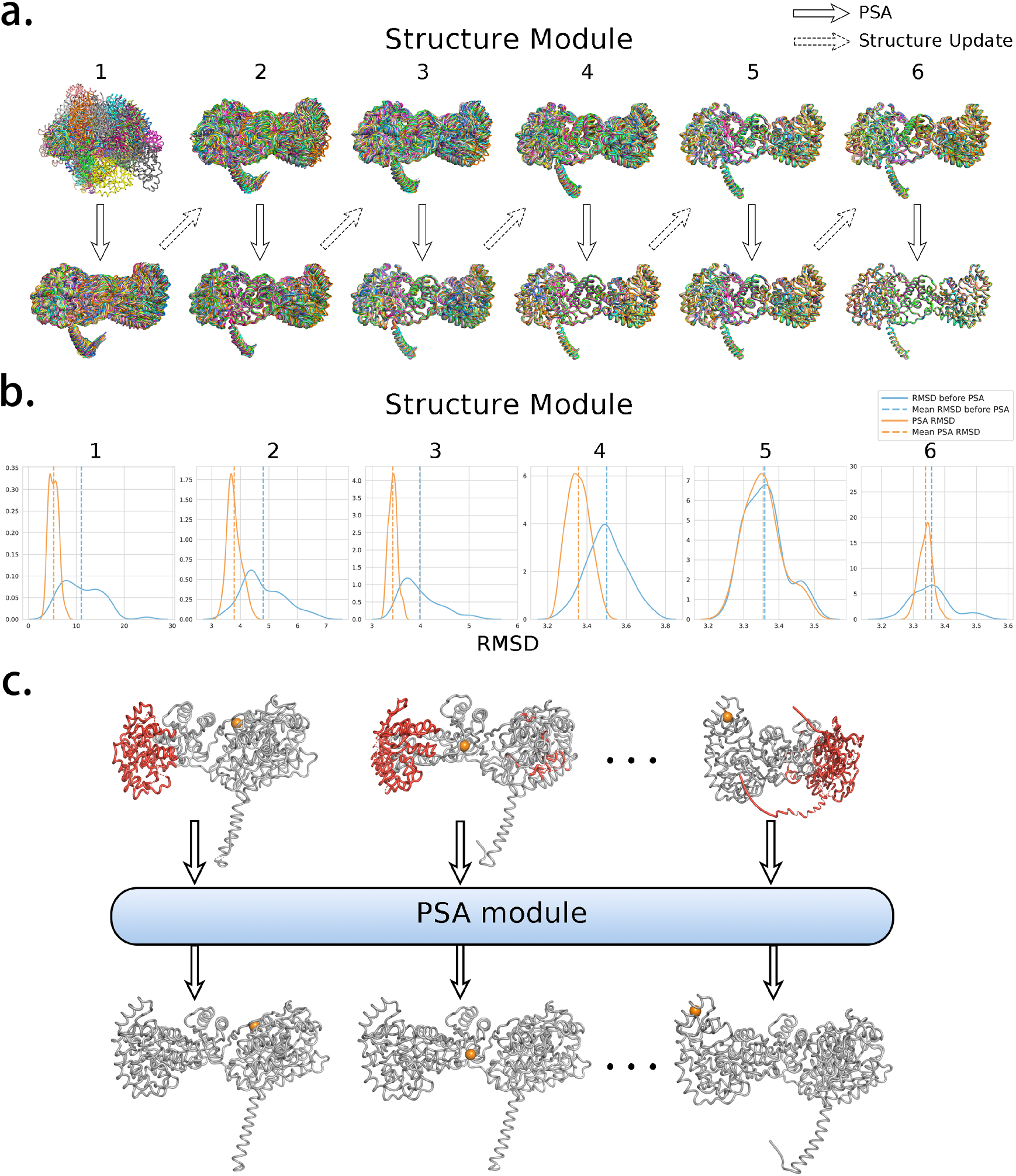
Case study on 8EE0_A by Cerebra. **a)** The folding process through 6 structure generation motifs in the Structure Module. Solid arrows represent the PSA processing within the structure generation motif, while dashed arrows represent the iterative coordinate updates between sequentially connected structure generation motifs. **b)** The probability density distributions of the RMSD values (against the ground truth structure) for the structures generated before (colored blue) and after (colored orange) PSA processing in each of the 6 structure generation motifs. **c)** Exemplar protein structures generated by the 1^*st*^ structure generation motif in the Structure Module. The first and second row show the structures before and after PSA processing, respectively, with poorly predicted regions highlighted in red. Structures in the same column present atomic coordinates obtained in the reference frame of the same anchor residue, as highlighted by orange spheres.

### PSA improves the prediction on quaternion

Similar to the translation prediction, in order to validate the role of PSA module in the rotation prediction, we collected the quaternion predictions Q_*ij*|*k*_ and the attention weights *ω*_Q,*k*_ along various rotation paths (see Equation 3) generated by each structure generation motif of the Structure Module for each residue pair *i* and *j* on the CAMEO targets. We then calculated the mean quaternion angular error, 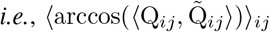 where 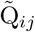 denotes the ground-truth quaternion and the inner product is calculated between the predicted and ground-truth unit quaternions. Again, Q_*ij*_ was derived by three different approaches: PSA attention-weighted Lie-algebra (Log–Exp) interpolation (see Equation 3), unweighted Lie-algebra (Log–Exp) interpolation, and the raw prediction Q_*ij*|*k*=*i*_. As shown in Table 3, PSA processing universally attains the lowest quaternion prediction error, outperforming both unweighted interpolation and the raw prediction. Interestingly, averaging over the top1/3/5 paths with the highest overall attention weights could achieve nearly the same level of low prediction errors as the full PSA processing (see Table S2), which implies that through model training, the PSA module effectively places focuses on a few important paths for information integration.

**Table 3.** Prediction errors in quaternion by different methods.

| Residue Distance $r_{ij}$ | Method | Structure Module | | | | | |
| --- | --- | --- | --- | --- | --- | --- | --- |
|  |  | 1 | 2 | 3 | 4 | 5 | 6 |
| [0, 10) | Raw | 0.339 | 0.326 | 0.323 | 0.315 | 0.294 | 0.291 |
|  | Avg | 0.328 | 0.314 | 0.280 | 0.278 | 0.236 | 0.231 |
|  | PSA | **0.301** | **0.253** | **0.244** | **0.234** | **0.226** | **0.219** |
| [10, 20) | Raw | 0.355 | 0.344 | 0.338 | 0.316 | 0.315 | 0.305 |
|  | Avg | 0.370 | 0.323 | 0.317 | 0.315 | 0.286 | 0.283 |
|  | PSA | **0.310** | **0.308** | **0.290** | **0.286** | **0.279** | **0.273** |
| [20, 30) | Raw | 0.403 | 0.393 | 0.389 | 0.366 | 0.361 | 0.350 |
|  | Avg | 0.416 | 0.371 | 0.362 | 0.360 | 0.333 | 0.330 |
|  | PSA | **0.356** | **0.352** | **0.337** | **0.332** | **0.327** | **0.321** |
| [30, $+\infty$) | Raw | 0.492 | 0.490 | 0.489 | 0.467 | 0.451 | 0.442 |
|  | Avg | 0.497 | 0.460 | 0.453 | 0.446 | 0.421 | 0.418 |
|  | PSA | **0.440** | **0.438** | **0.426** | **0.418** | **0.413** | **0.408** |
| [0, $+\infty$) | Raw | 0.405 | 0.397 | 0.393 | 0.370 | 0.367 | 0.355 |
|  | Avg | 0.414 | 0.370 | 0.364 | 0.361 | 0.331 | 0.328 |
|  | PSA | **0.356** | **0.354** | **0.335** | **0.329** | **0.324** | **0.319** |

For each type of classification, the method showing the lowest prediction error is highlighted in bold. The prediction error is the quaternion angular difference in radians, calculated as the arccosine of the inner product between the predicted and ground-truth unit quaternions. Raw denotes the raw prediction, Avg denotes unweighted Lie-algebra (Log-Exp) interpolation, and PSA denotes attention-weighted Lie-algebra (Log-Exp) interpolation. $r_{ij}$ is the distance between residue $i$ and residue $j$ in the native protein structure.

We also evaluated the PCC between the PSA attention weights and the quaternion path metrics. As shown in Table 4, a universal negative correlation is still present, indicating that PSA tends to rely on intermediate residues located in-between the target residue and the reference anchor residue for quaternion inference. However, magnitudes of PCC values are smaller than the corresponding values in the translation analysis (see Table 2), consistent with our intuition that the orientational coupling is generally weaker than the positional coupling between two distantly placed objects.

**Table 4.** Pearson correlation between quaternion path metrics and PSA attention weights Path Metric Residue Distance *r*_*ij*_ Structure Module.

| Path Metric | Residue Distance $r_{ij}$ | Structure Module | | | | | |
| --- | --- | --- | --- | --- | --- | --- | --- |
|  |  | 1 | 2 | 3 | 4 | 5 | 6 |
| Length | [0, 10) | -0.119 | -0.141 | -0.167 | -0.161 | -0.178 | -0.123 |
|  | [10, 20) | -0.164 | -0.201 | -0.185 | -0.166 | -0.201 | -0.168 |
|  | [20, 30) | -0.151 | -0.188 | -0.161 | -0.142 | -0.177 | -0.168 |
| | [30, $+\infty$) | -0.094 | -0.123 | -0.121 | -0.096 | -0.116 | -0.104 |
| | [0, $+\infty$) | -0.121 | -0.153 | -0.153 | -0.132 | -0.157 | -0.128 |
| Angle | [0, 10) | -0.167 | -0.212 | -0.352 | -0.283 | -0.277 | -0.027 |
|  | [10, 20) | -0.242 | -0.318 | -0.417 | -0.288 | -0.301 | -0.074 |
|  | [20, 30) | -0.198 | -0.255 | -0.293 | -0.208 | -0.224 | -0.117 |
| | [30, $+\infty$) | -0.114 | -0.152 | -0.170 | -0.113 | -0.126 | -0.091 |
| | [0, $+\infty$) | -0.163 | -0.218 | -0.273 | -0.192 | -0.203 | -0.080 |

Length refers to the absolute path length in the Euclidean space, while Angle means the cosine value of the scalar angle of the path. $r_{ij}$ is the distance between residue $i$ and residue $j$ in the native protein structure. Direct paths between $i$ and $j$ (*i.e.* when $k = i$ ) are excluded from the evaluation of the Angle metric to avoid the lack of angle definition.

### Cerebra effectively assists hallucination-based *de novo* protein backbone design

To investigate whether the inference efficiency of Cerebra could facilitate iterative protein design, we repurposed Cerebra for hallucination-based *de novo* backbone generation by optimizing random amino acid sequences to maximize predicted confidence. Here, we developed two protocols: the refined protocol employs a gradual 5,000-step search, whereas the fast protocol preferentially mutates low-confidence regions and thus reduces the optimization to 1,500 steps. Under the length-dependent production setting, the mean retained pLDDT increases from an initial range of 0.253–0.431 to 0.704–0.862 after optimization. The refined protocol provides additional confidence gains for shorter proteins, whereas the fast protocol offers a more favorable balance between confidence and computational cost for longer proteins (Figure S9). Hence, we adopted the refined protocol for small proteins of 50–100 residues and the fast protocol for large proteins of 150–200 residues in the following evaluation.

We next evaluated the designability of generated backbones using ProteinMPNN sequence design followed by ESMFold refolding. For each backbone, the lowest backbone RMSD among 10 independently designed and refolded sequences was used for evaluation. Among the 400 evaluated backbones, 98.5% achieve a minimum backbone RMSD below 2.0 Å, with the success rate of 99%, 100%, 99% and 96% for proteins of 50, 100, 150 and 200 residues, respectively (Figure 6a). The median of minimum RMSDs ranges from 0.539 to 0.833 Å, demonstrating consistently high designability across different protein lengths. Besides the folding confidence in pLDDT by ESMFold (Figure 6b), we also evaluated the structural novelty of generated protein backbones by searching against structures in the PDB database using Foldseek (Figure 6c). The median of top-hit PDB-TM scores (*i*.*e*. the highest TM-score among PDB structures) range from 0.712 to 0.801 across the four sequence lengths. Among the 385 backbones with valid Foldseek results, 34.8% have the PDB-TM score below 0.7 and 10.6% have the score below 0.6, indicating that the generated backbones cover a broad range of structural dissimilarities to known PDB proteins.

**Figure 6.**
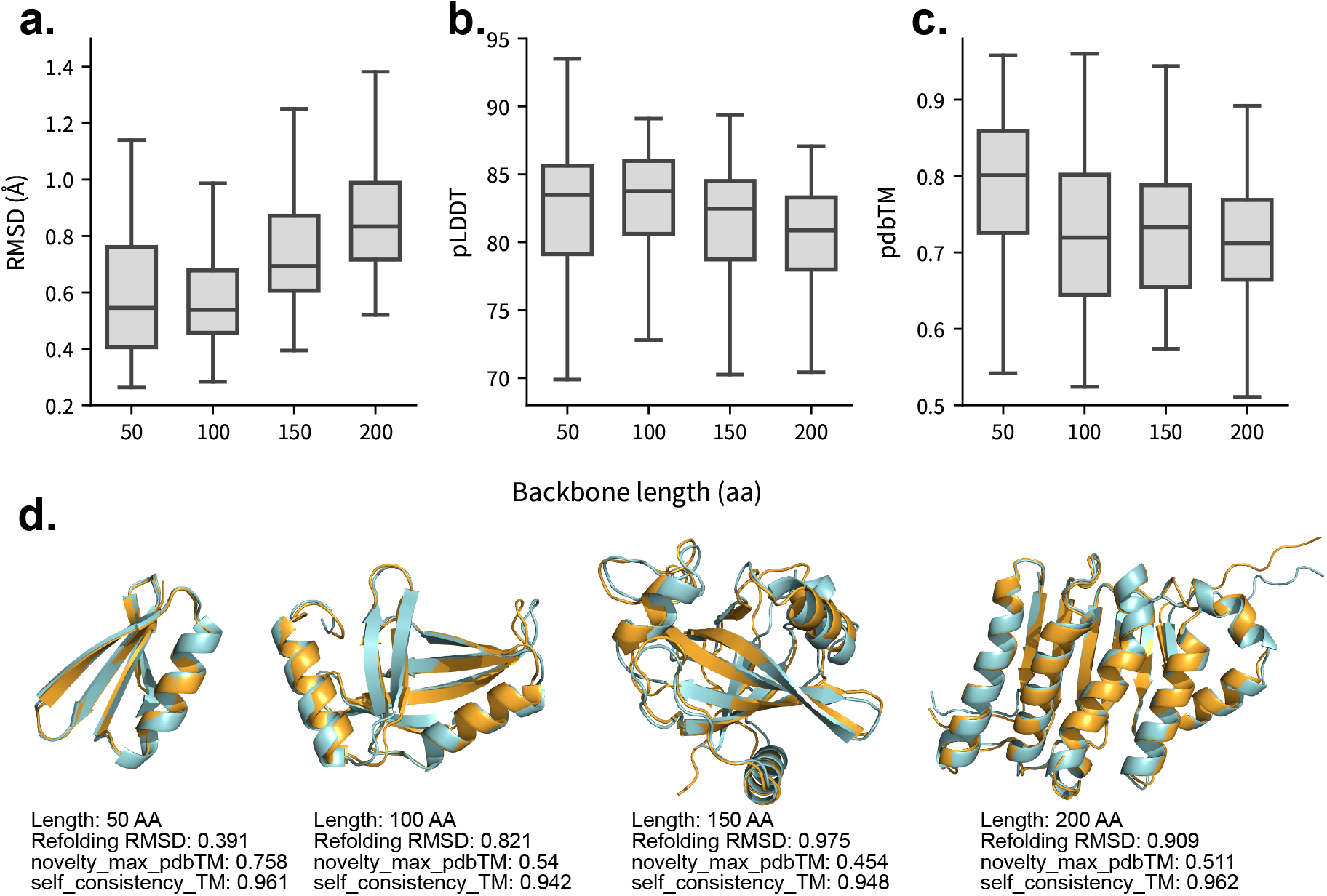
Representative examples of Cerebra-generated protein backbones via hallucination. **a)** Minimum backbone RMSD among 10 ProteinMPNN–ESMFold refolded structures for each generated backbone. **b)** ESMFold pLDDT corresponding to the minimum-RMSD structure. **c)** PDB-TM score of the top Foldseek hit against the PDB. Each length group contains 100 backbones, with valid Foldseek results available for 90, 100, 99 and 96 backbones, respectively. In panels (**a**)–(**c**), center lines and numerical labels indicate medians, boxes represent interquartile ranges, and whiskers extend to 1.5 times the interquartile range, while outliers are not shown. **d)** Structural alignments of representative designs containing 50, 100, 150 and 200 residues. Cerebra-generated backbones are colored orange, and ESMFold-predicted structures of ProteinMPNN-designed sequences are colored blue. Protein length, refolding RMSD, maximum PDB-TM score, and self-consistency TM-score are reported below each example. Smaller PDB-TM scores indicate lower structural similarity to known protein structures in the PDB.

In Figure 6d, we present four successful protein designs comprising 50, 100, 150 and 200 residues, respectively. The refolding RMSDs of these designed sequences are 0.391, 0.821, 0.975, and 0.909 Å, respectively, with the self-consistency TM-score ranging from 0.942 to 0.962. Moreover, the PDB-TM scores range from 0.454 to 0.758, demonstrating that the high designability is achievable across backbones with different levels of novelty to known structures. Together, these results confirm that the inference efficiency of Cerebra enables effective exploration of sequence space and generation of highly designable protein backbones, including a novel subset with low structural similarity to proteins in the PDB.

## Discussion

In this work, we developed a novel framework Cerebra for the task of accurate protein structure prediction. Particularly, the Cerebra model adopts a brand new Structure Module, which utilizes the 2D information to generate multiple sets of atomic coordinates in parallel and then relies on an innovative attention-based PSA module to further suppress the structure prediction errors by effectively integrating the complementary information among the multiple sets of predicted structures. Our approach remarkably enhances the computational efficiency, achieving an acceleration of at least 6 folds in model training, based on benchmark testing against the state-of-the-art OpenFold model. The current Cerebra model was trained on limited computational resources, but achieves a comparable performance to the published version of the OpenFold model on benchmark testing set. Considering its outstanding behavior in improving the computational efficiency and in reducing the training consumption, the current Cerebra model has already provided a good starting point for ordinary research groups. Specifically, an independent research group could either retrain or fine-tune this model using freely chosen datasets, aiming to address special biological problems. Moreover, the superiority of Cerebra in model inference time supports its application in scenarios where intensive model calling is required, for instance, in the hallucination-based *de novo* protein design, as evidenced in this study.

Many computational methods adopt the graph representation to describe a protein, where the nodes correspond to amino acid residues and the edges quantify inter-residue interactions. The graph representation of proteins achieves successes in many practical applications, including protein structure prediction^13,14^, protein functional analysis^38–41^, protein-ligand interaction^42–44^, etc., due to the adequate consideration of the pairwise residue-residue interaction in addition to the 1D sequence information. Historically, theories or models purely based on pairwise descriptions have confronted great challenges, since they are unable to provide a perfect description for the complex physical, chemical or biological systems, particularly when the three-body and higher-order many-body effects are salient. Clearly, the three-body effects are usually non-negligible in proteins, manifested by the allosteric regulations on protein functions. Previous graph-based models partially circumvent this problem by stacking multiple graph neural network (GNN) or graph transformer layers, since an arbitrary pair of nodes located in neighboring layers could communicate through a common intermediate node, thus implicitly composing triangular interactions. Obviously, explicit inclusion of three-body description within the design of neural networks will provide a more efficient solution. In this respect, the triangular self-attention and triangular multiplicative update in the AlphaFold2 design are the best-known representatives, both of which are reported to make significant contributions to the state-of-the-art performance of AlphaFold2. Unfortunately, such triangular modules can only handle the scalar variables due to the lack of consideration on the geometric relationship between vectors or tensors in the Cartesian coordinate system during information integration. In this work, we propose the PSA attention module to explicitly handle three-body effects without disrupting the intrinsic geometric relationship between the translation vectors and rotation matrices of residues under processing. In the current implementation, PSA can adjust the poorly predicted coordinates by leveraging the coordinate information of one additional relay residue located in-between the target residue and the reference anchor, and the consistency loss can further suppress geometric incompatibility after the PSA adjustment. When treating more complicated systems in the future, paths composed of two relay residues could be constructed to enable the processing of geometric relationship among four residues (*i*.*e*. four-body effects) in exactly the same manner. Hence, the innovative PSA attention developed in this work provides a new solution to graph-based models, for properly handling the many-body effects in the three-dimensional Euclidean space.

## Data availability

Input features for all proteins tested in this work and their evaluation results are available at https://zenodo.org/records/21698980.

## Code availability

The model of Cerebra is implemented in PyTorch^45^. All codes, including the training scripts, are freely downloadable at https://github.com/Gonglab-THU/Cerebra. The optimized model parameters are available at https://zenodo.org/records/21698980. An online server for Cerebra is available at https://structpred.life.tsinghua.edu.cn/server_cerebra.html.

## Accessibility statement

Cerebra is freely accessible to all academic users under the MIT License. A freely accessible web server is available at https://structpred.life.tsinghua.edu.cn/server_cerebra.html.

## Author contribution

H. G. proposed the methodology and designed the experiment. J. H. and H. G. made the initial design for the network architecture. J. H. and W. W. implemented the coding and preliminary model training. W. W made local modification on the architecture and finalized the overall model training. B. Z. performed the hallucination-based protein design. Y. C. contributed to the model benchmarking and evaluation. J. H., W. W., B. Z. and Y. C. analyzed the results. H. G. supervised the whole process. J. H., W. W. and H. G. wrote the initial draft, and B. Z. contributed to manuscript revision. All authors agreed with the final manuscript.

## Acknowledgments

This work has been supported by the National Natural Science Foundation of China (#32671644 & #32171243), by the Ministry of Science and Technology of China (#2023YFF1204400) and by the Beijing Frontier Research Center for Biological Structure.

## References

1. Jumper, J., Evans, R., Pritzel, A., et al. Highly accurate protein structure prediction with AlphaFold. Nature 596, 583–589. ISSN: 1476-4687. 10.1038/s41586-021-03819-2 (2021).

2. Gao, M., Nakajima An, D., Parks, J. M., et al. AF2Complex predicts direct physical interactions in multimeric proteins with deep learning. Nature Communications 13. ISSN: 2041-1723. 10.1038/s41467-022-29394-2 (2022).

3. Bryant, P., Pozzati, G. & Elofsson, A. Improved prediction of protein-protein interactions using AlphaFold2. Nature Communications 13. ISSN: 2041-1723. 10.1038/s41467-022-28865-w (2022).

4. Evans, R., O’Neill, M., Pritzel, A., et al. Protein complex prediction with AlphaFold-Multimer. 10.1101/2021.10.04.463034 (2021).

5. Wong, F., Krishnan, A., Zheng, E. J., et al. Benchmarking AlphaFold-enabled molecular docking predictions for antibiotic discovery. Molecular Systems Biology 18. ISSN: 1744-4292. 10.15252/msb.202211081 (2022).

6. Hekkelman, M. L., de Vries, I., Joosten, R. P., et al. AlphaFill: enriching AlphaFold models with ligands and cofactors. Nature Methods 20, 205–213. ISSN: 1548-7105. 10.1038/s41592-022-01685-y (2022).

7. Tsaban, T., Varga, J. K., Avraham, O., et al. Harnessing protein folding neural networks for peptide–protein docking. Nature Communications 13, 176. ISSN: 2041-1723. 10.1038/s41467-021-27838-9 (2022).

8. Akdel, M., Pires, D. E. V., Pardo, E. P., et al. A structural biology community assessment of AlphaFold2 applications. Nature Structural & Molecular Biology 29, 1056–1067. ISSN: 1545-9985. 10.1038/s41594-022-00849-w (2022).

9. Pak, M. A., Markhieva, K. A., Novikova, M. S., et al. Using AlphaFold to predict the impact of single mutations on protein stability and function. PLOS ONE 18, 1–9. 10.1371/journal.pone.0282689 (2023).

10. Hu, M., Yuan, F., Yang, K., et al. Exploring evolution-aware & -free protein language models as protein function predictors in Advances in Neural Information Processing Systems (eds Koyejo, S., Mohamed, S., Agarwal, A., et al.) 35 (Curran Associates, Inc., 2022), 38873–38884. https://proceedings.neurips.cc/paper_files/paper/2022/file/fe066022bab2a6c6a3c57032a1623c70-Paper-Conference.pdf.

11. Zhang, Y., Li, P., Pan, F., et al. Applications of AlphaFold beyond Protein Structure Prediction. BioRxiv. 10.1101/2021.11.03.467194 (2021).

12. Cheng, J., Novati, G., Pan, J., et al. Accurate proteome-wide missense variant effect prediction with AlphaMissense. Science 381. ISSN: 1095-9203. 10.1126/science.adg7492 (2023).

13. Lin, Z., Akin, H., Rao, R., et al. Evolutionary-scale prediction of atomic-level protein structure with a language model. Science 379, 1123–1130. https://www.science.org/doi/abs/10.1126/science.ade2574 (2023).

14. Wu, R., Ding, F., Wang, R., et al. High-resolution de novo structure prediction from primary sequence. bioRxiv. https://www.biorxiv.org/content/early/2022/07/22/2022.07.21.500999 (2022).

15. Fang, X., Wang, F., Liu, L., et al. A method for multiple-sequence-alignment-free protein structure prediction using a protein language model. Nature Machine Intelligence 5, 1087–1096. ISSN: 2522-5839. 10.1038/s42256-023-00721-6 (2023).

16. Yim, J., Trippe, B. L., De Bortoli, V., et al. SE(3) diffusion model with application to protein backbone generation in Proceedings of the 40th International Conference on Machine Learning (eds Krause, A., Brunskill, E., Cho, K., et al.) 202 (PMLR, 2023), 40001–40039. https://proceedings.mlr.press/v202/yim23a.html.

17. Lin, Y. & AlQuraishi, M. Generating Novel, Designable, and Diverse Protein Structures by Equivariantly Diffusing Oriented Residue Clouds 2023. https://arxiv.org/abs/2301.12485.

18. Wicky, B. I. M., Milles, L. F., Courbet, A., et al. Hallucinating symmetric protein assemblies. Science 378, 56– 61. https://www.science.org/doi/abs/10.1126/science.add1964 (2022).

19. Frank, C., Khoshouei, A., Fuß, L., et al. Scalable protein design using optimization in a relaxed sequence space. Science 386, 439–445. https://www.science.org/doi/abs/10.1126/science.adq1741 (2024).

20. Pacesa, M., Nickel, L., Schellhaas, C., et al. One-shot design of functional protein binders with BindCraft. Nature 646, 483–492. ISSN: 1476-4687. 10.1038/s41586-025-09429-6 (2025).

21. Baek, M., Anishchenko, I., Humphreys, I. R., et al. Efficient and accurate prediction of protein structure using RoseTTAFold2. 10.1101/2023.05.24.542179 (2023).

22. Baek, M., McHugh, R., Anishchenko, I., et al. Accurate prediction of protein–nucleic acid complexes using RoseTTAFoldNA. Nature Methods 21, 117–121. ISSN: 1548-7105. 10.1038/s41592-023-02086-5 (2023).

23. Krishna, R., Wang, J., Ahern, W., et al. Generalized biomolecular modeling and design with RoseTTAFold All-Atom. Science 384, eadl2528. https://www.science.org/doi/abs/10.1126/science.adl2528 (2024).

24. Watson, J. L., Juergens, D., Bennett, N. R., et al. De novo design of protein structure and function with RFdiffusion. Nature 620, 1089–1100. ISSN: 1476-4687. 10.1038/s41586-023-06415-8 (2023).

25. Abramson, J., Adler, J., Dunger, J., et al. Accurate structure prediction of biomolecular interactions with AlphaFold 3. Nature. ISSN: 1476-4687. 10.1038/s41586-024-07487-w (2024).

26. Wohlwend, J., Corso, G., Passaro, S., et al. Boltz-1: Democratizing Biomolecular Interaction Modeling. bioRxiv. 10.1101/2024.11.19.624167 (2024).

27. Passaro, S., Corso, G., Wohlwend, J., et al. Boltz-2: Towards Accurate and Efficient Binding Affinity Prediction. bioRxiv. 10.1101/2025.06.14.659707 (2025).

28. Chai Discovery. Chai-1: Decoding the molecular interactions of life. bioRxiv. 10.1101/2024.10.10.615955 (2024).

29. Chai Discovery. Zero-shot antibody design in a 24-well plate. bioRxiv. 10.1101/2025.07.05.663018 (2025).

30. ByteDance AML AI4Science Team, Chen, X., Zhang, Y., et al. Protenix: Advancing Structure Prediction Through a Comprehensive AlphaFold3 Reproduction. bioRxiv. 10.1101/2025.01.08.631967 (2025).

31. Ahdritz, G., Bouatta, N., Floristean, C., et al. OpenFold: retraining AlphaFold2 yields new insights into its learning mechanisms and capacity for generalization. Nature Methods. ISSN: 1548-7105. 10.1038/s41592-024-02272-z (2024).

32. OpenFold Consortium. OpenFold3 Training Data Registry of Open Data on AWS. Accessed: 2026-03-31. 2025. https://registry.opendata.aws/openfold3/.

33. Mirdita, M., Schütze, K., Moriwaki, Y., et al. ColabFold: making protein folding accessible to all. Nature Methods 19, 679–682. ISSN: 1548-7105. 10.1038/s41592-022-01488-1 (2022).

34. Zhang, Y. & Skolnick, J. Scoring function for automated assessment of protein structure template quality. Proteins: Structure, Function, and Bioinformatics 57, 702–710. 10.1002/prot.20264 (2004).

35. Mariani, V., Biasini, M., Barbato, A., et al. lDDT: a local superposition-free score for comparing protein structures and models using distance difference tests. Bioinformatics 29, 2722–2728. 10.1093/bioinformatics/btt473 (2013).

36. Dauparas, J., Anishchenko, I., Bennett, N., et al. Robust deep learning-based protein sequence design using ProteinMPNN. Science 378, 49–56. 10.1126/science.add2187 (2022).

37. Van Kempen, M., Kim, S. S., Tumescheit, C., et al. Fast and accurate protein structure search with Foldseek. Nature Biotechnology 42, 243–246. 10.1038/s41587-023-01773-0 (2024).

38. Gligorijević, V., Renfrew, P. D., Kosciolek, T., et al. Structure-based protein function prediction using graph convolutional networks. Nature Communications 12. ISSN: 2041-1723. 10.1038/s41467-021-23303-9 (2021).

39. Chen, J., Qian, Y., Huang, Z., et al. Enhancing Protein Solubility Prediction through Pre-trained Language Models and Graph Convolutional Neural Networks in 2023 IEEE International Conference on Bioinformatics and Biomedicine (BIBM) (IEEE, 2023). 10.1109/BIBM58861.2023.10385858.

40. Zhai, H., Hou, H., Luo, J., et al. DGDTA: dynamic graph attention network for predicting drug–target binding affinity. BMC Bioinformatics 24. ISSN: 1471-2105. 10.1186/s12859-023-05497-5 (2023).

41. Chen, L., Wu, R., Zhou, F., et al. HybridGCN for protein solubility prediction with adaptive weighting of multiple features. Journal of Cheminformatics 15. ISSN: 1758-2946. 10.1186/s13321-023-00788-8 (2023).

42. Xia, Y., Xia, C.-Q., Pan, X., et al. GraphBind: protein structural context embedded rules learned by hierarchical graph neural networks for recognizing nucleic-acid-binding residues. Nucleic Acids Research 49, e51–e51. ISSN: 0305-1048. 10.1093/nar/gkab044 (2021).

43. Fang, Y., Jiang, Y., Wei, L., et al. DeepProSite: structure-aware protein binding site prediction using ESMFold and pretrained language model. Bioinformatics 39 (ed Cowen, L.) ISSN: 1367-4811. 10.1093/bioinformatics/btad718 (2023).

44. Yuan, Q., Chen, S., Rao, J., et al. AlphaFold2-aware protein–DNA binding site prediction using graph transformer. Briefings in Bioinformatics 23. ISSN: 1477-4054. 10.1093/bib/bbab564 (2022).

45. Paszke, A., Gross, S., Massa, F., et al. in Proceedings of the 33rd International Conference on Neural Information Processing Systems (Curran Associates Inc., Red Hook, NY, USA, 2019). https://dl.acm.org/doi/10.5555/3454287.3455008.

