## Supplementary Information for "Cerebra: a computationally efficient framework for accurate protein structure prediction"

### 1 Supplementary Results

#### 1.1 Performance on the CAMEO 2024 validation set

During model development, the cleaned CAMEO 2024 dataset, comprising 643 protein targets, was used exclusively for model selection, including checkpoint and quaternion-interpolation selection. No gradient-based model training or parameter updating were performed using these targets. Candidate Cerebra checkpoints were evaluated on this validation set, and the five checkpoints with the highest mean TM-scores were selected for subsequent evaluation. The CAMEO 2025 dataset was not involved in model selection and was retained as the held-out test set reported in **Results**.

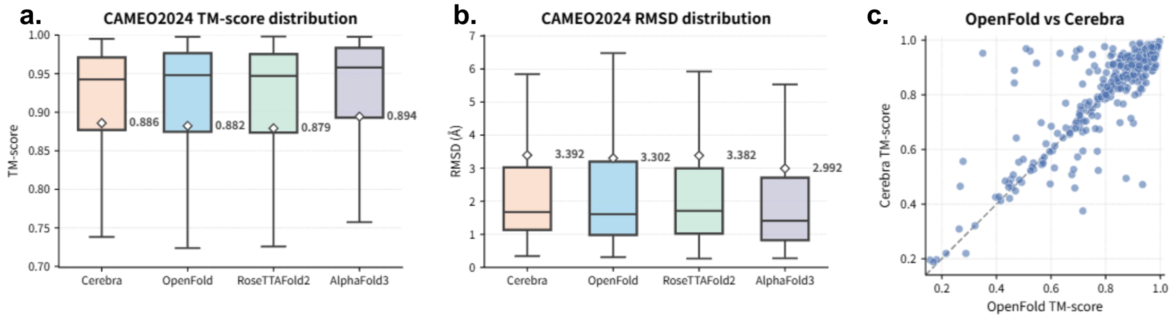


**Figure S1. Comparison of protein structure prediction performance on the CAMEO 2024 set. a–b)** Distributions of TM-score (**a**) and RMSD (**b**) for structures predicted by OpenFold, Cerebra, RoseTTAFold2, and AlphaFold3 on 643 CAMEO targets. Center lines represent medians, boxes indicate interquartile ranges, and whiskers extend to the most extreme values within 1.5 times the interquartile range. White diamonds and adjacent values indicate means. **c)** Per-target comparison of TM-scores obtained by Cerebra and OpenFold. The dashed line indicates equal performance.

Using the selected checkpoints, Cerebra achieves a mean TM-score of 0.886 on the CAMEO 2024 validation set, in comparison with 0.882 by OpenFold. Cerebra obtains a higher TM-score than OpenFold on 252 out of the 643 targets, including 24 targets for which its TM-score advantage exceeds 0.1 (Figure S1). These results characterize the validation performance used for model selection and are consistent with the subsequent results obtained on the independent CAMEO 2025 test set (Figure 2).

#### 1.2 The depths of Evoformer and Structure Module

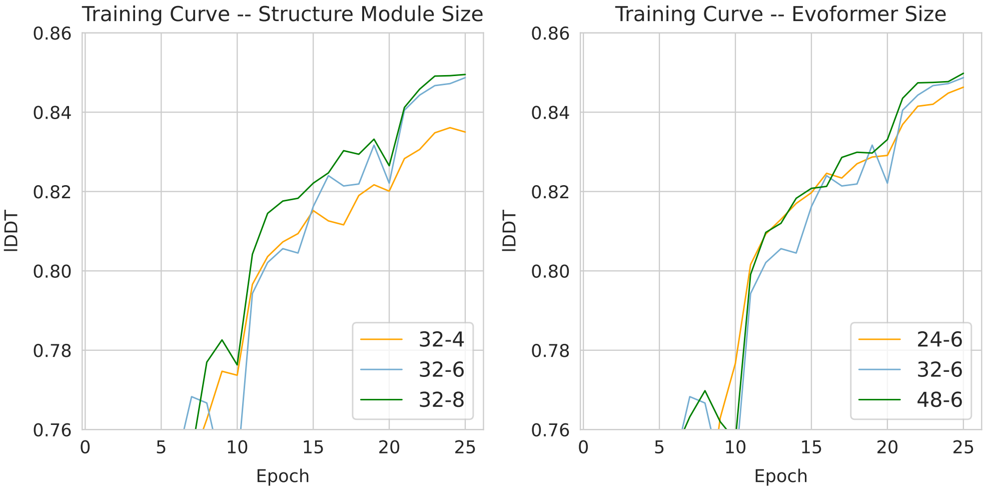


**Figure S2. The training curves for the models with different depths of Evoformer and Structure Module.** In the legend, the labels in the format of "$m$-$n$" denote the network depth, where $m$ is the number of Evoformer motifs and $n$ is the number of Structure Module motifs. Hence, 32-6 refers to the standard Cerebra architecture of 32 Evoformer motifs and 6 Structure Module motifs.

We trained Cerebra models with different numbers of Evoformer motifs and Structure Module motifs on the 10K dataset to explore the effects of network depth on the model performance (Figure S2). Clearly, increasing either the number of Evoformer motifs or that of Structure Module motifs will elicit improvement in the model performance. In this study, we adopt a medium level of network depth (32 Evoformer motifs and 6 Structure Module motifs in comparison to the numbers of 48 and 8 in AlphaFold2) to allow model training on a single GPU.

##### 1.3 Topics for anchor residues

###### 1.3.1 The number of anchor residues during training

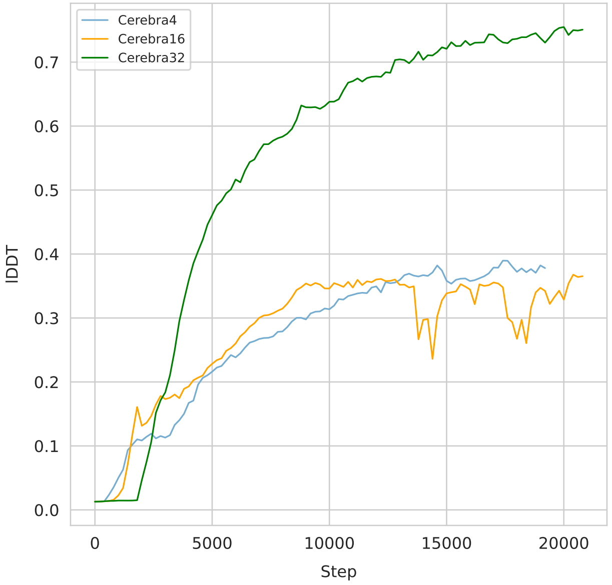


**Figure S3. The training curves of the models with different numbers of anchor residues.**

In the initial training process, the number of anchor residues is an important hyper-parameter that needs optimization. Therefore, in addition to the default setting of 32 anchor residues, we also trained Cerebra models with fewer anchor residues to investigate the possibility of further reducing the GPU memory cost. Unfortunately, Cerebra models with 4 or 16 anchor residues show significantly weakened performance (Figure S3). Specifically, the training lDDT lingers below 0.4 even after 20,000 training steps, remarkably lower than the lDDT score above 0.7 in the standard Cerebra model with 32 anchor residues. Introduction of more anchor residues may bring further improvement, particularly for large proteins, but at the cost of increased GPU memory consumption. Consequently, we retained the 32 anchor residues in the standard Cerebra model.

##### 1.3.2 The number of anchor residues during inference

Due to the variability in protein size in practical prediction, the number of anchors indeed influences the quality of predicted structures. We evaluated the Cerebra model on varying anchor numbers for CAMEO 2024 and CAMEO 2025 targets grouped by protein length. For each target, the TM-score was first averaged across all evaluated anchor-number settings. Targets with a mean TM-score below 0.2 were regarded as complete prediction failures and excluded. The same retained target set was then used for all anchor-number settings and grouped by protein length. The results are shown in Table S1.

**Table S1. The number of anchor residues during inference**

| Length | < 96 | | [96, 224) | | [224, 324] | | (324, 500] | | > 500 | |
|---|---|---|---|---|---|---|---|---|---|---|
| Anchor Number | TM-score | lDDT | TM-score | lDDT | TM-score | lDDT | TM-score | lDDT | TM-score | lDDT |
| 12 | 0.731 | 0.816 | 0.854 | 0.848 | 0.874 | 0.861 | 0.908 | 0.860 | 0.855 | 0.825 |
| 24 | 0.734 | 0.818 | 0.854 | 0.850 | 0.873 | 0.862 | 0.910 | 0.865 | 0.856 | 0.831 |
| 32 | 0.731 | 0.817 | 0.856 | 0.852 | 0.875 | 0.863 | 0.910 | 0.867 | 0.857 | 0.832 |
| 48 | 0.730 | 0.815 | 0.856 | 0.852 | 0.875 | 0.864 | 0.909 | 0.867 | 0.856 | 0.834 |
| 56 | 0.730 | 0.815 | 0.855 | 0.852 | 0.871 | 0.862 | 0.908 | 0.863 | 0.857 | 0.834 |

In general, the effect of anchor number is modest, without clear monotonic performance increase with the number of anchors. Across the five length groups, the range between the highest and lowest values is at most 0.004 for TM-score and at most 0.009 for lDDT. Because of the exclusion of targets with TM-score below 0.2, these values characterize the sensitivity to the anchor number among successfully predicted targets rather than the overall performance on the complete CAMEO datasets.

For proteins shorter than 96 residues, 24 anchors produces the highest TM-score and lDDT values of 0.734 and 0.818, respectively, whereas additional anchors provide no benefit. For proteins of 96–324 residues, 32 and 48 anchors generally achieve the highest or near-highest scores. For proteins of 324–500 residues, 32 anchors reach the highest TM-score of 0.910, tying with 48 anchors for the highest lDDT of 0.867. The clearest benefit of using more anchors is observed for proteins of at least 500 residues, for which increasing the anchor number from 12 to 48 improves lDDT from 0.825 to 0.834, while TM-score changes only slightly.

Because additional anchors increase memory usage, the results indicate that 24 anchors are sufficient for short proteins, 32 anchors provide a favorable accuracy–efficiency trade-off for proteins of 96–500 residues, and increasing the number to 48 is beneficial mainly for proteins of at least 500 residues. This analysis was conducted post hoc to characterize sensitivity to the number of anchors; the inference settings used for the primary CAMEO 2025 evaluation were fixed beforehand.

##### 1.3.3 Schemes for anchor residue selection

Cerebra by default tends to distribute anchor residues relatively evenly across the entire protein sequence, ensuring that each residue is located in the neighborhood of an anchor residue. Here, we try an alternative approach, that is, evenly distributing the anchor residues spatially upon the protein structure. The basic protocol is designed as follows:

1) The default sequence-based anchor selection scheme is used in the first recycling iteration (cycle = 1), due to the lack of knowledge on the protein structure.
2) In each of the subsequent recycling iterations, anchor residues are selected based on the protein structure predicted from the preceding recycling iteration. Specifically, all residues are first clustered based on the inter-residue distance matrix (calculated from the protein structure) using the Kmeans clustering algorithm of Scikit-learn[1] (version 0.23.2), and the residues closest to the centroids of clusters are chosen as anchor residues, which jointly compose the AnchorList for the protein structure prediction in the current recycling iteration.

We evaluated this structure-based anchor selection scheme on the CAMEO dataset. The average TM-score and lDDT are 0.839 and 0.802, respectively, slightly lower than the respective values of 0.842 and 0.816 obtained by the default sequence-based anchor selection scheme. Pairwise comparison of these two schemes, as shown in Figure S4, negates the presence of significant difference between them.

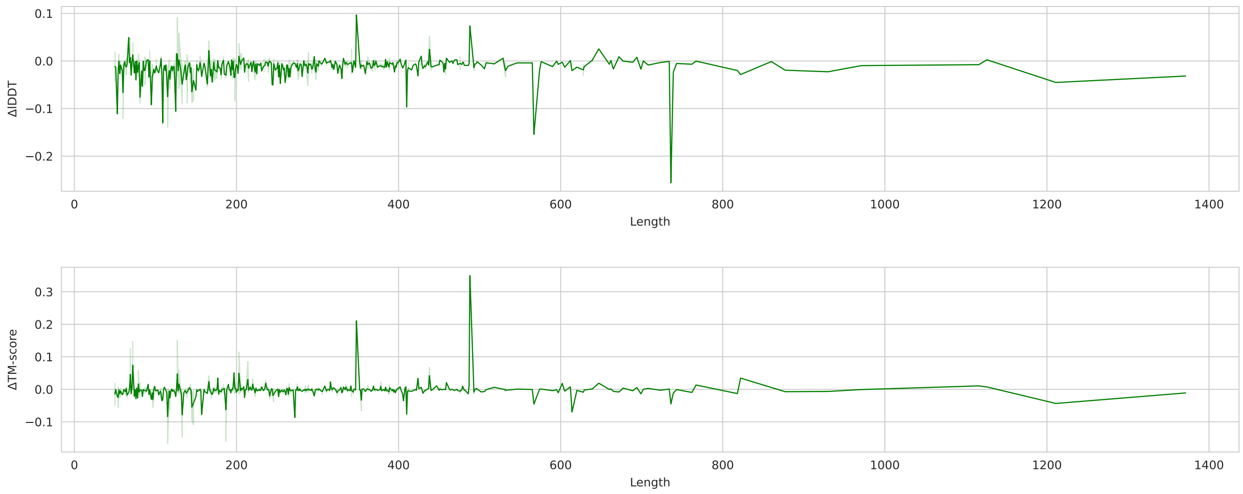


**Figure S4. Evaluation of structure-based vs. sequence-based anchor selection schemes on the CAMEO dataset.** Here, the difference in lDDT and TM-score between the two schemes are plotted against the protein length for all targets in the CAMEO set.

We speculate the following potential reasons for the trivial contribution of the structure-based anchor selection scheme:

1) The PSA module in Cerebra is designed to leverage multiple "synthesized" paths for coordinate updating. Hence, as long as the number of anchor residues is sufficient, each residue could find an anchor residue in its vicinity, no matter whether the anchor residues are distributed based on sequence or structure. Moreover, the PSA module will automatically assign higher weights to the paths mediated by relay residues spatially located between the source and target residues.
2) Due to the lack of prior knowledge about the ground-truth protein structure, which is also the problem we are trying to solve, spatial clustering of residues can be only performed based on the protein structure predicted

from the preceding recycling iteration. Hence, errors in the predicted structure itself may hinder correct spatial clustering, a deficit that makes the sequence-based scheme a more robust solution.

#### 1.4 Topics for quaternion prediction

##### 1.4.1 Quaternion prediction using the top paths assigned by PSA

**Table S2. Errors in the quaternion prediction by PSA and top paths**

| **Residue Distance $r_{ij}$** | **Method** | **Structure Module** | | | | | |
|---|---|---|---|---|---|---|---|
| | | 1 | 2 | 3 | 4 | 5 | 6 |
| [0, 10) | PSA | **0.301** | **0.253** | **0.244** | **0.234** | **0.226** | **0.219** |
| | Top1 | 0.305 | 0.254 | **0.244** | 0.236 | 0.230 | 0.222 |
| | Top3 | 0.303 | 0.259 | 0.247 | 0.235 | 0.227 | 0.220 |
| | Top5 | 0.303 | 0.254 | 0.246 | **0.234** | 0.227 | **0.219** |
| [10, 20) | PSA | **0.310** | **0.308** | **0.290** | **0.286** | **0.279** | **0.273** |
| | Top1 | 0.321 | 0.311 | 0.292 | 0.287 | 0.282 | 0.276 |
| | Top3 | 0.313 | 0.309 | 0.291 | 0.287 | 0.280 | 0.274 |
| | Top5 | 0.312 | 0.309 | 0.291 | **0.286** | 0.280 | 0.274 |
| [20, 30) | PSA | **0.356** | **0.352** | **0.337** | **0.332** | **0.327** | **0.321** |
| | Top1 | 0.364 | 0.355 | 0.338 | **0.332** | 0.330 | 0.324 |
| | Top3 | 0.360 | 0.353 | 0.339 | 0.333 | 0.328 | 0.322 |
| | Top5 | 0.358 | 0.353 | 0.338 | **0.332** | 0.328 | 0.322 |
| [30, $+\infty$) | PSA | **0.440** | **0.438** | **0.426** | **0.418** | **0.413** | **0.408** |
| | Top1 | 0.444 | 0.440 | 0.427 | **0.418** | 0.415 | 0.410 |
| | Top3 | 0.442 | 0.439 | 0.428 | 0.419 | **0.413** | 0.409 |
| | Top5 | **0.440** | 0.439 | 0.427 | 0.419 | **0.413** | 0.409 |
| [0, $+\infty$) | PSA | **0.356** | **0.354** | **0.335** | **0.329** | **0.324** | **0.319** |
| | Top1 | 0.361 | 0.358 | 0.337 | 0.331 | 0.327 | 0.321 |
| | Top3 | 0.357 | 0.356 | 0.337 | 0.331 | 0.325 | **0.319** |
| | Top5 | **0.356** | **0.354** | 0.336 | 0.330 | 0.325 | **0.319** |

For each type of classification, the method showing the lowest prediction errors is highlighted in bold. The prediction error is the angular difference in radians between the prediction and the label. PSA stands for the attention-based weighted average as designed in Cerebra, while Top1/3/5 denote averaging over the top1/3/5 paths with the highest overall attention weights. $r_{ij}$ is the distance between residue $i$ and residue $j$ in the native protein structure.

Clearly, as shown in Table S2, when using the top paths with the highest attention weights, the errors of quaternion prediction are close to PSA processing, for residue pairs located at all ranges and among all structure generation motifs in the Structure Module.

##### 1.4.2 Comparison of quaternion interpolation methods

**Table S3. Comparison of prediction errors by different quaternion-interpolation methods**

| Method | | Structure Module | | | | | |
|---|---|---|---|---|---|---|---|
| | | 1 | 2 | 3 | 4 | 5 | 6 |
| Nlerp | Raw | 0.415 | 0.393 | 0.393 | 0.385 | 0.378 | 0.376 |
| | Avg | 0.395 | 0.376 | 0.372 | 0.363 | 0.357 | 0.357 |
| | PSA | 0.367 | 0.361 | 0.353 | 0.349 | 0.346 | 0.346 |
| Markley | Avg | 0.388 | 0.367 | 0.362 | 0.357 | 0.353 | 0.349 |
| | PSA | 0.382 | 0.361 | 0.356 | 0.352 | 0.348 | 0.344 |
| Log-Exp | Avg | 0.351 | 0.349 | 0.346 | 0.344 | 0.343 | 0.344 |
| | PSA | 0.349 | 0.347 | 0.344 | 0.342 | 0.342 | 0.342 |

Raw denotes the raw prediction results, Avg refers to the simple unweighted averaging, while PSA stands for the attention-based weighted average as designed in Cerebra.

Cerebra used normalized linear interpolation (Nlerp) in the quaternion PSA throughout initial and large-scale training. Before fine-tuning, we used the resulting checkpoint to compare Nlerp, Markley quaternion averaging, and Lie algebra (Log–Exp) on the CAMEO 2024 validation set, with all other model parameters fixed. Nlerp averages quaternion components directly and requires normalization; Markley averaging uses an outer-product construction invariant to the antipodal representations Q and $-$Q, whereas Lie algebra (Log–Exp) combines relative rotations in the tangent space.

As shown in Table S3, Lie algebra (Log–Exp) produces the lowest quaternion prediction errors for both simple unweighted averaging and PSA attention-weighted interpolation across the six Structure Module motifs. It also achieves the highest lDDT and TM-score values throughout the Structure Module (Figure S5). Based on this comparison, we selected Lie-algebra interpolation for the subsequent fine-tuning stage and retained it as the quaternion interpolation method in the PSA module of the final Cerebra model. Its detailed implementation is provided in Algorithm 4 (see Supplementary Information 3.3 for details).

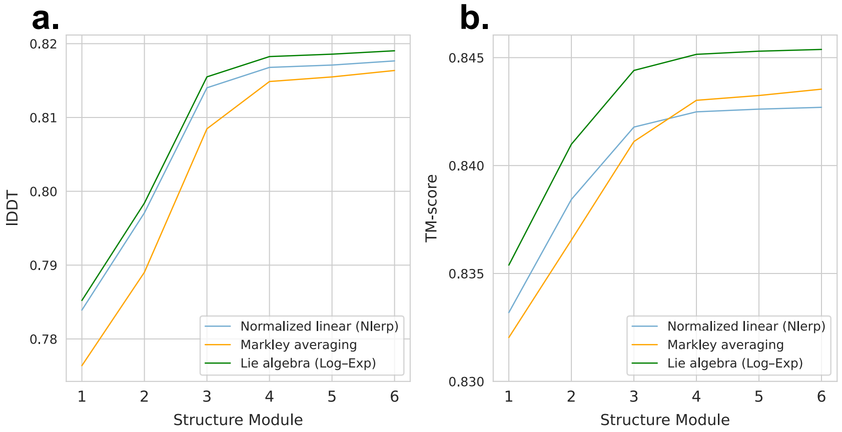


**Figure S5. The impact of quaternion interpolation methods on model performance.** All three methods were evaluated on the CAMEO 2024 validation set using the Nlerp-trained checkpoint obtained before fine-tuning. Panels show lDDT (**a**) and TM-score (**b**) across the six structure generation motifs of the Structure Module.

#### 1.5 About model training efficiency

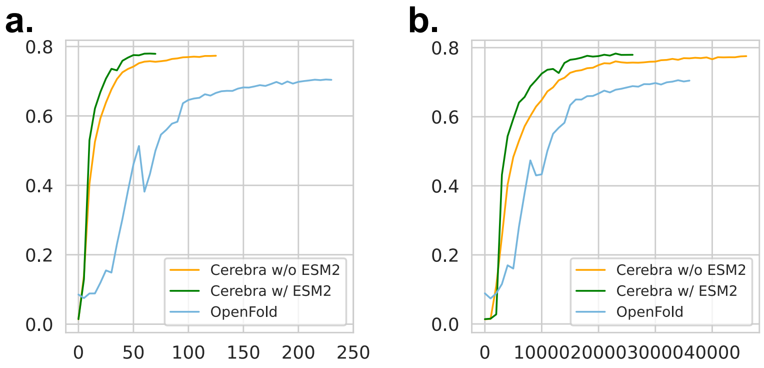


**Figure S6. Training curves of Cerebra and OpenFold showing the growth of model performance during training.** The model performance is represented by lDDT in the vertical axis, whereas the training process is represented in the horizontal axis by GPU hours (**a**) and training steps (**b**), respectively.

We compared the training efficiency of Cerebra and OpenFold on the same 10K dataset using identical MSA inputs and structural labels. To isolate the contribution of the model architecture, ESM-2 embeddings were disabled in the primary Cerebra setting. Training efficiency was evaluated according to both cumulative GPU hours and the number of parameter-update steps. Cerebra reached training lDDT values of 0.6 and 0.7 after 21 and 34 GPU hours, respectively, whereas OpenFold required 91 and 205 GPU hours to reach the same performance levels (Figure S6a). When pre-trained ESM-2 embeddings were additionally introduced, Cerebra reached the corresponding lDDT values after only 14 and 24 GPU hours. Cerebra also required approximately half the number of update steps needed by OpenFold to achieve comparable training performance (Figure S6b). These results demonstrate that the Cerebra architecture substantially accelerates model convergence and that pre-trained sequence embeddings provide a further improvement in training efficiency. Both models were trained separately on a single NVIDIA A100 GPU with 80 GB of memory.

#### 1.6 Topics for model inference

##### 1.6.1 Effects of recycling

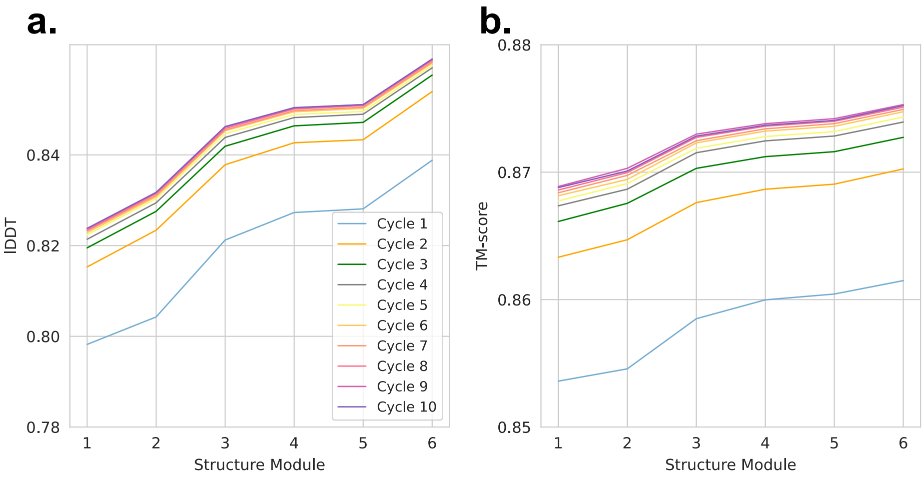


**Figure S7. The impact of recycling cycle numbers on the prediction accuracy.**

Figure S7 presents the changes of lDDT and TM-score along the successive structure generation motifs of the Structure Module, when a specific checkpoint of the Cerebra model is applied on the CAMEO test set for structure prediction using various numbers of recycling iterations. Clearly, the quality of predicted structures enhances both along the Structure Module motifs and along the recycling iterations during practical inference. For instance, in the first cycle, lDDT and TM-score start from merely 0.798 and 0.854, respectively, but improve to 0.839 and 0.861, respectively. Further iterative inference cycles markedly benefit the model's predictive accuracy. When focusing on the $6^{th}$ Structure Module motif, lDDT and TM-score values rise from 0.839 and 0.861 in the first cycle to 0.859 and 0.874 in the fourth cycle, and to 0.861 and 0.875 in the tenth cycle, respectively.

The evident enhancement in model prediction accuracy with successive recycling iterations can be attributed to the following reasons. Firstly, due to GPU memory constraints, Cerebra employs an MSA subsampling approach, incorporating only 128 homologous sequences per cycle as model inputs. Sampling different portions of the overall MSA across different cycles effectively utilizes more co-evolutionary information, thereby enriching the model's input. Secondly, recycling can be considered as a method to virtually increase model depth in the GPU-memory-constrained environment.

As shown in the Figure S7, the model's predictive performance tends to improve with increasing cycles. However, we opted for a default recycle setting of three cycles as the final output, aligning with the practice of the state-of-the-art models like AlphaFold2, which ensures a fair performance comparison. Our model is fully open-sourced, allowing users to modify the number of iterations to potentially achieve higher prediction accuracy.

##### 1.6.2 Potential for complex prediction

Similar to AlphaFold2, Cerebra is capable of predicting the structure of heteromeric protein complexes without the need for additional training. Unlike the standard monomeric protein structure prediction, feature preparation for the heteromeric protein complex structure prediction involves some modifications. Here are the detailed steps involved in the feature preparation process.

**MSA Search Process:**

The MSA search adopts the ColabFold protocol[2], which involves searching for the MSAs of each monomer and then concatenating these sequences.

**ESM-2 Embedding Process:**

1) The heteromeric protein sequences are first linked together using a sequence of 32 glycine (GLY) residues.
2) The concatenated sequence is then embedded using the ESM-2 model.
3) After obtaining the ESM-2 embedding, the inserted GLY segments are removed to obtain the embedding for the target protein sequences only.

**Complex Inference Process:**

The process for predicting the structure of the protein complex is similar to that used for monomeric proteins. We have applied this methodology to predict a few heteromeric protein complexes, as shown in Figure S8.

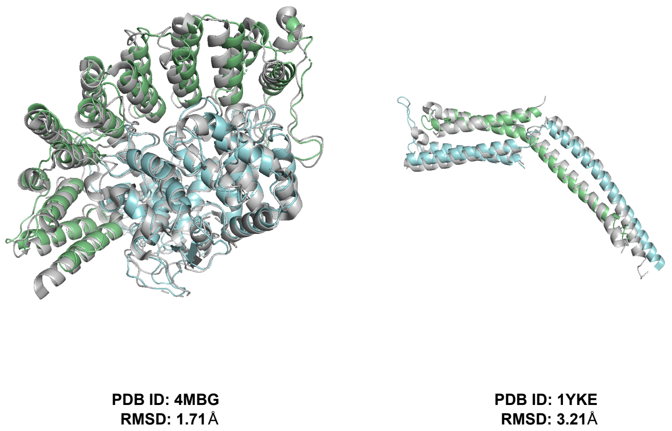


**Figure S8. Cerebra can predict the structure of protein complexes without retraining.** The native structure is shown in gray, whereas the predicted structure is colored in a chain-wise manner. In the dimeric structures shown here, the two chains are colored in green and cyan, respectively.

#### 1.7 Optimization of hallucination protocols

We compared the Cerebra-hallucination-fast and Cerebra-hallucination-refined protocols across protein lengths of 50, 100, 150 and 200 residues. The fast protocol used 1,500 sequence update steps and preferentially mutated low-confidence regions, whereas the refined protocol performed a more gradual search over 5,000 steps. Across the four lengths, the fast protocol increased the mean retained pLDDT from 0.253–0.415 to 0.704–0.785, while the refined protocol increased it from 0.273–0.431 to 0.749–0.862 (Figure S9). Although the refined protocol reached higher endpoint pLDDT values at all lengths, its advantage over the fast protocol decreased to 0.063 and 0.045 for proteins of 150 and 200 residues, respectively, despite requiring more than three times as many sequence updates. We therefore used the refined protocol for proteins of 50 and 100 residues and the fast protocol for proteins of 150 and 200 residues.

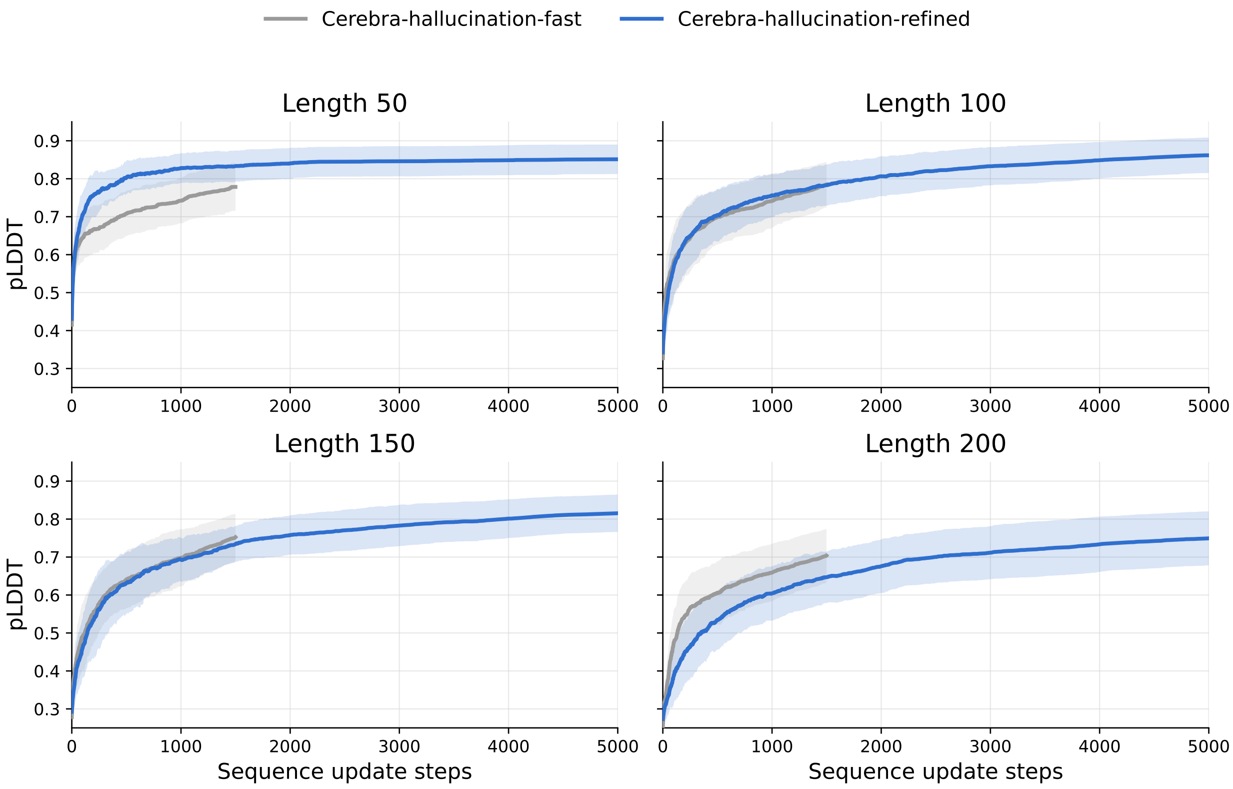


**Figure S9. Comparison of hallucination optimization protocols across protein lengths.** Mean retained pLDDT is plotted against the number of sequence update steps for proteins of 50, 100, 150 and 200 residues. Gray and blue curves represent the Cerebra-hallucination-fast and Cerebra-hallucination-refined protocols, respectively. Solid curves and shaded regions indicate the mean and $\pm 1$ standard deviation across 50 optimization trajectories.

#### 2 Feature and Notation

The input data of Cerebra can be classified into three main categories: ESM-2 embeddings of the target sequences, MSAs and sequence indices. The ESM-2 embeddings of the target sequences are generated using the ESM-2 (3B) model, while the MSAs and the sequence indices are the same as those used in AlphaFold2. We refer to the MSA as 1D information, the latent variables composed of residue pairs as 2D information, and the latent variables related to atomic coordinates as structure information (or 3D information). The structure information includes the translations (as vectors) and rotations (as quaternions) of residues in the reference coordinate system, denoted as T (for translation) and Q (for quaternion), respectively. In addition to the latent variables, the network also utilizes AnchorList, which is a list of anchor residues. The symbols and descriptions involved in the model are listed in Table S4.

**Table S4. Feature and notation**

| Short Name | Describe | Dimension |
|---|---|---|
| $K$ | Number of residues in AnchorList | 12/24/32 |
| $M$ | Number of sequences in MSA | 128 (training) / 256 (evaluation) |
| $L$ | Target sequence length | |
| $E_1$ | 1D feature dimension | 256 |
| $E_2$ | 2D feature dimension | 128 |
| | MSA feature (MSA Feat) | $[M, L, 46]$ |
| | Target sequence feature (Target Feat) | $[L, 21]$ |
| x1D | 1D feature (1D Feat) | $[M, L, E_1]$ |
| x2D | 2D feature (2D Feat) | $[E_2, L, L]$ |
| T | Translation | $[K, L, 3]$ |
| Q | Quaternion | $[K, L, 4]$ |

#### 3 Basic Formulas

##### 3.1 Local coordinate system

To calculate the relative translation and rotation between residues, a local coordinate system has to be established for each residue. We use a plane formed by the three atoms $C_\alpha$, C, and N as the $xy$ plane of the coordinate system. The $C_\alpha$ atom is located at the origin, the $C_\alpha \to$ C vector represents the $x$-axis, and the component of $C_\alpha \to$ N orthogonal to the $x$-axis represents the $y$-axis. The normal vector of the $xy$ plane represents the $z$-axis. Following this definition, the rotation matrix of the local coordinate system in the global frame can be estimated by Algorithm 1.

**Algorithm 1** Rotation matrix

def BackboneRotation($\mathbf{N}$, $\mathbf{C}$, $\mathbf{C}_\alpha \in R^3$):

1: $\mathbf{v}_1$ = $\mathbf{C}$ - $\mathbf{C}_\alpha$
2: $\mathbf{v}_2$ = $\mathbf{N}$ - $\mathbf{C}_\alpha$
3: $\mathbf{e}_1$ = $\mathbf{v}_1 / \|\mathbf{v}_1\|$
4: $\mathbf{e}_2$ = $\mathbf{v}_2 / \|\mathbf{v}_2\|$
5: $\mathbf{e}_2$ = $\mathbf{e}_2 - (\mathbf{e}_1 \cdot \mathbf{e}_2)\mathbf{e}_1$
6: $\mathbf{e}_2$ = $\mathbf{e}_2 / \|\mathbf{e}_2\|$
7: $\mathbf{e}_3$ = $\mathbf{e}_1 \times \mathbf{e}_2$
8: $\mathbf{R}$ = concat($\mathbf{e}_1$, $\mathbf{e}_2$, $\mathbf{e}_3$)

##### 3.2 Quaternion basic formulas

In our model, we use quaternions to describe rotation matrices of residues. Therefore, it is necessary to briefly describe the properties of quaternions and the operations involved.

Quaternions are mathematical entities that extend the concept of complex numbers to four dimensions, consisting of a scalar part and a vector part. Here, we use Q to represent quaternions, where $w$ and $\mathbf{v}$ denote the real part and the imaginary part of the quaternion, respectively.

$$\mathrm{Q} = [w, i, j, k] = [w, \mathbf{v}] \quad \text{(E1)}$$

Quaternion composition, which represents the composition of two rotations, can be expressed by Formula E2. In this study, we will use "$\odot$" to represent the composition of quaternions.

$$\mathrm{Q}_1 \odot \mathrm{Q}_2 = [w_1, \mathbf{v}_1][w_2, \mathbf{v}_2] = [w_1 * w_2 - \mathbf{v}_1 \cdot \mathbf{v}_2, \mathbf{v}_2 \times \mathbf{v}_1 + w_2 * \mathbf{v}_1 + w_1 * \mathbf{v}_2] \quad \text{(E2)}$$

Negating quaternion values results in inconsistent rotation paths in space, but the rotation angles remain the same:

$$\mathrm{Q} = \mathrm{Q} * [-1, -1, -1, -1] \quad \text{(E3)}$$

Negating the imaginary part of a quaternion represents a rotation in the opposite direction:

$$\mathrm{Q}^{-1} = \mathrm{Q} * [1, -1, -1, -1] \quad \text{(E4)}$$

Formula E5 and Formula E6 express the inter-conversion relationship between quaternions and rotation matrices:

$$\mathbf{R} = \begin{pmatrix} w^2+i^2-j^2-k^2 & 2ij-2wk & 2jw+2ik \\ 2ij+2kw & w^2-i^2+j^2-k^2 & 2jk-2wi \\ 2ik-2wj & 2jk+2wi & w^2-i^2-j^2+k^2 \end{pmatrix} \tag{E5}$$

$$\begin{cases} w = \text{sqrt}(\text{trace}(\mathbf{R})+1)/2 \\ i = (\mathrm{R}_{2,1}-\mathrm{R}_{1,2})/(4\times w) \\ j = (\mathrm{R}_{0,2}-\mathrm{R}_{2,0})/(4\times w) \\ k = (\mathrm{R}_{1,0}-\mathrm{R}_{0,1})/(4\times w) \end{cases} \tag{E6}$$

A quaternion, except for its magnitude, can be converted into a normalized quaternion (Algorithm 2). In addition, this study defines that the real part of a quaternion is non-negative.

**Algorithm 2** Norm Quaternion

def NormQuaternion($\mathrm{Q} \in R^4$):

1: $\mathrm{Q} = \mathrm{Q}/\|\mathrm{Q}\|$
2: $\mathrm{Q} = \text{sign}(\text{sign}(\mathrm{Q}[0])+0.5) * \mathrm{Q}$

If we consider coordinates as the imaginary part of a quaternion and define its real part as 0, then quaternions can be combined with coordinates. Algorithm 3 demonstrates the values of the coordinates after undergoing quaternion transformation.

**Algorithm 3** Translation By Quaternion

def TranslationByQuaternion($\mathrm{T} \in R^3$, $\mathrm{Q} \in R^4$):

1: $\mathrm{Q} = \mathrm{Q}/\|\mathrm{Q}\|$
2: $\mathrm{T}_4 = [0, \mathrm{T}]$
3: $\mathrm{T} = \mathrm{Q} \odot \mathrm{T}_4 \odot \mathrm{Q}^{-1}$
4: $\mathrm{T} = \mathrm{T}[1:]$

##### 3.3 Averaging quaternions

In this study, the rotation of residue $j$ in the local coordinate system of residue $i$ can be represented by $\mathrm{Q}_{ij}$. Given a rotation path mediated by an intermediate residue $k$, $\mathrm{Q}_{ij|k}$ can be calculated as $\mathrm{Q}_{ik} \odot \mathrm{Q}_{kj}$. Given $K$ available paths and their normalized PSA attention weights $\omega_{\mathrm{Q},k}$, Cerebra combines these rotations in the tangent space rather than averaging their quaternion components directly (Algorithm 4).

For each source anchor residue $i$, the implementation selects the corresponding diagonal element of the path tensor as the base quaternion. This element is the direct path $\mathrm{Q}_{ij|i} = \mathrm{Q}_{ij}$, obtained when the intermediate anchor residue is identical to the source anchor. Because Q and $-\mathrm{Q}$ represent the same rotation, each path quaternion is first aligned to the hemisphere of this base quaternion:

$$\widetilde{\mathrm{Q}}_{ij|k} = s_k \mathrm{Q}_{ij|k}, \qquad s_k = \begin{cases} 1, & \langle \mathrm{Q}_{ij|k}, \mathrm{Q}_{ij|i} \rangle \geq 0, \\ -1, & \langle \mathrm{Q}_{ij|k}, \mathrm{Q}_{ij|i} \rangle < 0. \end{cases} \tag{E7}$$

For a unit quaternion $\mathrm{Q} = [w, \mathbf{v}]$ with $\theta = 2\arccos(w)$, the logarithmic map used in the implementation is:

$$\mathrm{Log}(\mathrm{Q}) = \alpha(w)\mathbf{v}, \qquad \alpha(w) = \begin{cases} \dfrac{\theta}{\sin(\theta/2)}, & \sin(\theta/2) \geq \epsilon, \\ 2 + \dfrac{1-w^2}{3}, & \sin(\theta/2) < \epsilon. \end{cases} \tag{E8}$$

Conversely, for a rotation vector $\mathrm{V} \in \mathbb{R}^3$ with $\theta = \|\mathrm{V}\|_2$, the exponential map is:

$$\mathrm{Exp}(\mathrm{V}) = [\cos(\theta/2), \beta(\theta)\mathrm{V}], \qquad \beta(\theta) = \begin{cases} \dfrac{\sin(\theta/2)}{\theta}, & \theta \geq \epsilon, \\ \dfrac{1}{2} - \dfrac{\theta^2}{48}, & \theta < \epsilon. \end{cases} \tag{E9}$$

Each aligned path rotation is expressed relative to the base quaternion, mapped to the Lie algebra, and combined linearly using the PSA attention weights. Following the quaternion-composition convention in Equation E2, the implementation is expressed as:

$$\begin{cases} \mathrm{V}_{ij|k} = \mathrm{Log}\left(\widetilde{\mathrm{Q}}_{ij|k} \odot \mathrm{Q}_{ij|i}^{-1}\right), \\ \overline{\mathrm{V}}_{ij} = \sum_{k=1}^{K} \omega_{\mathrm{Q},k}\mathrm{V}_{ij|k}, \\ \mathrm{Q}_{ij} = \mathrm{Norm}\left[\mathrm{Exp}(\overline{\mathrm{V}}_{ij}) \odot \mathrm{Q}_{ij|i}\right]. \end{cases} \tag{E10}$$

Small-angle Taylor approximations are included in both maps to avoid numerical instability. The input quaternion is normalized, the scalar input to $\arccos$ is clipped to $[-1+\epsilon, 1-\epsilon]$, and the final quaternion is normalized.

**Algorithm 4** Lie-algebra interpolation of path rotations

def LieAlgebraInterpolation($\mathrm{Q} \in [K, 4]$, $\omega \in [K]$):

1: $\mathrm{Q}_{\text{base}} \leftarrow \mathrm{Q}[a]$
2: $\mathrm{Q} \leftarrow \mathrm{where}(\langle \mathrm{Q}, \mathrm{Q}_{\text{base}} \rangle < 0, -\mathrm{Q}, \mathrm{Q})$
3: $\Delta\mathrm{Q} \leftarrow \mathrm{Q} \odot \mathrm{Q}_{\text{base}}^{-1}$
4: $\mathrm{V} \leftarrow \mathrm{Log}(\Delta\mathrm{Q})$
5: $\overline{\mathrm{V}} \leftarrow \sum_{k=1}^{K} \omega_k \mathrm{V}_k$
6: $\mathrm{Q}_{\text{weighted}} \leftarrow \mathrm{NormQuaternion}(\mathrm{Exp}(\overline{\mathrm{V}}) \odot \mathrm{Q}_{\text{base}})$
7: **return** $\mathrm{Q}_{\text{weighted}}$

The interpolation is applied independently to each prediction head. Before the head-wise outputs are averaged, they are sign-aligned to the quaternion predicted by the first head and then normalized, matching the implementation in the Path Synthesis module.

To compare the effects of different quaternion interpolation methods on protein structure prediction, we additionally implemented normalized linear interpolation (Nlerp) and Markley quaternion averaging as alternative approaches to the Lie-algebra interpolation used in the final Cerebra model. Nlerp is implemented as a direct weighted sum of the path quaternions followed by normalization:

$$\mathrm{Q}_{\text{weighted}} = \mathrm{Norm}(\sum_k \mathrm{Norm}(\mathrm{Q}_k) * w_k) \tag{E11}$$

Markley Algorithm (Algorithm 5) is a high-precision quaternion interpolation algorithm for weighted averaging of quaternions. Basically, it involves performing a singular value decomposition (SVD) on the weighted average of the quaternion covariance matrices, and the eigenvector corresponding to the largest eigenvalue represents the maximum likelihood estimate of the weighted average of the quaternions.

**Algorithm 5** Markley Averaging Quaternions

def MarkleyAveragingQuaternions(Q $\in [K, 4]$, $w \in [K]$):

1: $\mathbf{A} = \sum_{i}^{K} w_i * \text{outer}(\mathrm{Q}_i, \mathrm{Q}_i)$

2: $\mathrm{eigenValues}, \mathrm{eigenVectors} = \mathrm{SVD}(\mathbf{A})$

3: $\mathrm{Q} = \mathrm{eigenVectors}[\mathrm{eigenValues.argmax}]$

4: $\mathrm{Q}_{\mathrm{weighted}} = \mathrm{real}(\mathrm{Q})$

### 4 Cerebra Architecture

#### 4.1 Embedding Module and Evoformer Module

Our model consists of three modules: the Embedding Module, the Evoformer Module and the Structure Module. As shown in Figure S10, the input data for the Embedding Module consist of two parts. The first part includes the MSA features, target sequence features, ESM-2 embeddings and residue indices for the current recycling iteration. The second part contains the data obtained from the preceding recycling iteration, including the 1D features, the 2D features and the distance matrices calculated based on the multiple sets of atomic coordinates.

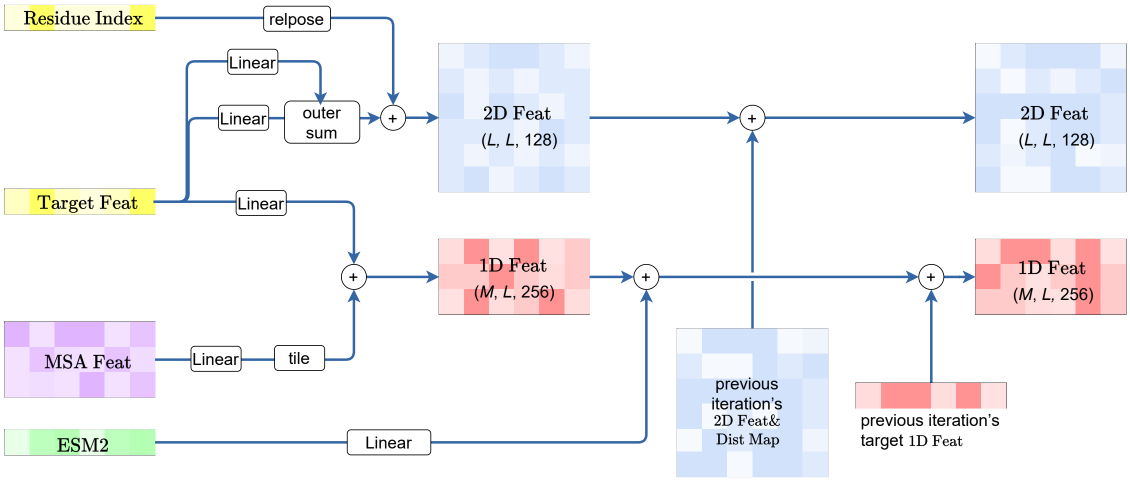


**Figure S10. Schematic architecture of the Embedding Module.**

The Embedding Module is set up in a generally similar way to AlphaFold2, with two main differences: 1) The template-related embedding module and additional MSA embedding module have been removed; 2) The ESM-2 embeddings need to be linearly projected to 256 dimensions and be added to each sequence:

$$\mathrm{m}_i = \mathrm{m}_i + \mathrm{Linear}(\mathrm{ESM2}). \tag{E12}$$

The Evoformer Module utilizes the OpenFold open-source code, with the number of motifs reduced from 48 to 32.

#### 4.2 Structure Module

In this study, we have designed a brand new Structure Module that can convert the 1D and 2D information output by the Evoformer Module into multiple sets of protein atomic coordinates. The overall architecture of each structure generation motif of the Structure Module is shown in Figure 1b, which includes a 1D encoder and a 1D decoder for the interaction between 1D features and 2D features, a structure encoder that integrates 2D features with structural information, a structure decoder that predicts new structures, and a PSA module that recombines multiple sets of coordinates. The pseudocode for the overall Structure Module is shown in Algorithm 6.

**Algorithm 6** StructureModule

**def** StructureModule(
  AnchorList $\in [Batch, K]$
  T $\in [Batch, K, L, 3]$=[..., [0, 0, 0]] ,
  Q $\in [Batch, K, L, 4]$=[..., [1, 0, 0, 0]]
  x1D $\in [Batch, M, L, E_1]$,
  x2D $\in [Batch, E_2, L, L]$ ):

for **Structure** in [StructureGenerationMotif1, StructureGenerationMotif2, ..., StructureGenerationMotif6]:
  x2D = **Structure.**1DEncoder(x1D, x2D)
  x2D = **Structure.**StructureEncoder(x2D, T, Q)
  x1D = **Structure.**1DDecoder(x1D, x2D)
  NewT, NewQ = **Structure.**StructureDecoder(x2D, T, Q, AnchorList)
  NewT = T + NewT
  NewT, NewQ, T_consistence, Q_consistence = **Structure.**PSA(x2D, NewT, NewQ, AnchorList)
  if not StructureGenerationMotif1:
    NewQ = NormQuaternion((Q + NewQ)/2.)
  T, Q = NewT, NewQ
SideChain = SideChainModule(x2D)
plDDT = plDDTModule(x2D,T)
**return** x1D, x2D, T, Q, T_consistence, Q_consistence, SideChain, plDDT

##### 4.2.1 Encoder

The module is divided into two separate parts: the 1D encoder that integrates the self-attention maps of 1D features into 2D features by Algorithm 7 and the structure encoder that integrates the structure information into 2D features by Algorithm 8.

**Algorithm 7** 1DEncoder

**def** 1DEncoder(
  x1D $\in [Batch, M, L, E_1]$,
  x2D $\in [Batch, E_2, L, L]$):

Q = ReShape(Linear(x1D)) $[Batch, Head, M, L, E_1]$
K = ReShape(Linear(x1D)) $[Batch, Head, M, L, E_1]$
Atten = OutProductOnL(Q, K) $[Batch, Head, M, L, L]$
Atten = SymmetrizationOnL(Atten)
x2D′ = Cat(x2D, Atten[:, :, 0], $\frac{1}{M}\sum_i^M$ Atten[:, :, $i$])
x2D′ = Conv2d(Norm$\underbrace{(\mathrm{LeakyRelu}(\mathrm{Conv2d}(\mathrm{Norm}(\mathrm{x2D}'))))}_{\times 2}$ $[Batch, E_2, L, L]$
**return** x2D + x2D′

The input features for the structure encoder are 2D features and the structural information that consists of $K$ sets of $C_\alpha$ coordinates and $K$ sets of quaternion quantifying rotation matrices. The encoding module needs to transform the structural information and integrate it into the 2D features. The structural information can be transformed in the following four ways (Figure S11):

1. Conversion of $K$ sets of quaternions to $L$ sets of quaternions;
2. Conversion of $K$ sets of coordinates to $L$ sets of coordinates with additional transformations;
3. Representation of the distance matrices for $C_\alpha$ atoms;
4. Representation of the distance matrices for $C_\beta$ atoms.

These transformations allow the encoding module to incorporate the structural information into the overall architecture and facilitate the subsequent 1D feature update as well as new structure generation in the decoder module.

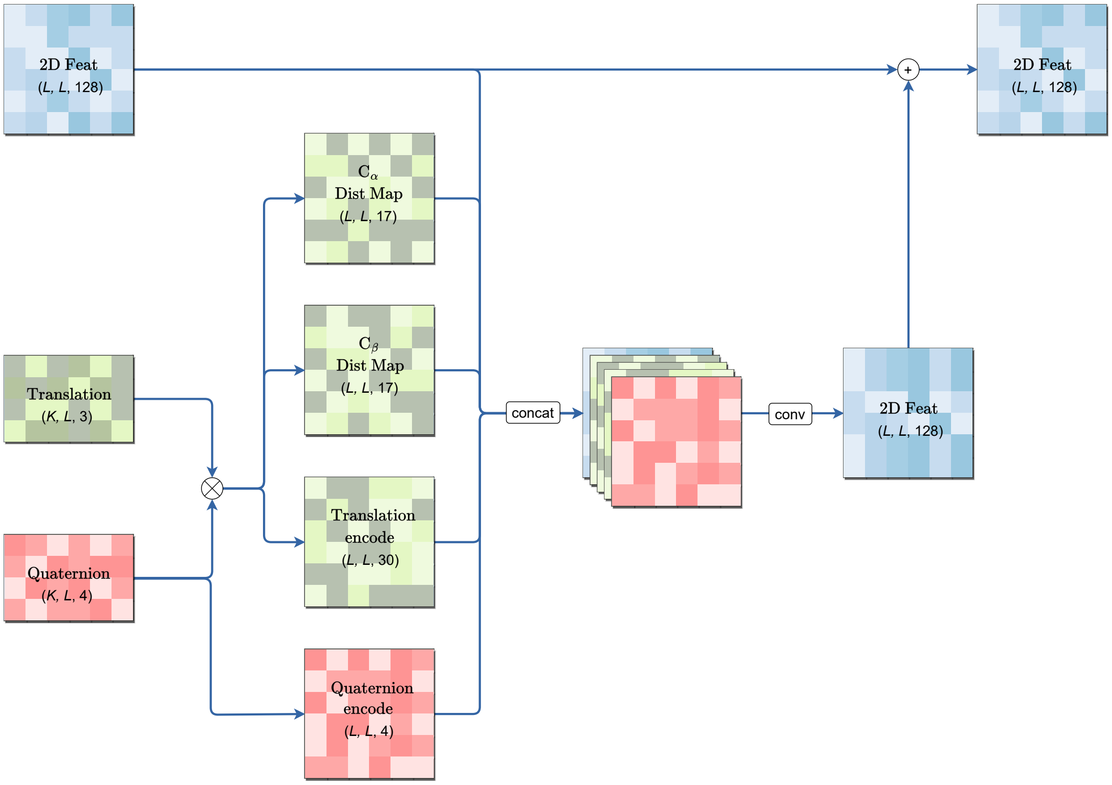


**Figure S11. Schematic architecture of the structure encoder.**

**Algorithm 8** StructureEncoder

**def** StructureEncoder(
    T $\in [Batch, K, L, 3]$),
    Q $\in [Batch, K, L, 4]$)
    x2D $\in [Batch, E_2, L, L]$ ):

**def** CombQT(x $\in [Batch, K, L, L, 1or3or4]$):
    $\text{x\_} = \text{Permute\&ReShape(x)}$ $[(Batch * K), 1or3or4, L, L]$
    $\text{Atten} = \text{Conv2d(Norm(LeakyRelu(Conv2d(Norm(x\_)))))}$ $[(Batch * K), 1, L, L]$
    $\text{Atten} = \text{Softmax(Permute\&ReShape(Atten))}$ $[Batch, L, L, 1, K]$
    $\text{x} = \text{Permute(x)}$ $[Batch, L, L, 1or3or4, K]$
    $\text{x} = \sum_i^K (\text{x}_i * \text{Atten}_i)$ $[Batch, L, L, 1or3or4]$
    **return** x
**def** Translation_K2L(T $\in [Batch, K, L, 3]$, Q $\in [Batch, K, L, 4]$):
    $\text{Q}_1 = \bar{\text{Q}}[:, :, :, \text{None}]$
    $\text{T}_1 = \text{T}[:, :, \text{None}, :] - \text{T}[:, :, :, \text{None}]$
    $\text{T}_L = \text{CombQT(TranslationRotation}(\text{T}_1, \text{Q}_1))$
    $\text{T}_L = \text{permute}(\text{T}_L)$ $[Batch, 3, L, L]$
    $\text{T}_{\text{encode}} = \text{T}_L * [1/10, 1/20, ..., 1/100]$ $[Batch, 30, L, L]$
    **return** $\text{T}_{\text{encode}}$
**def** Quaternion_K2L(Q $\in [Batch, K, L, 4]$):
    $\text{Q}_1 = \bar{\text{Q}}[:, :, :, \text{None}]$
    $\text{Q}_2 = \text{Q}[:, :, \text{None}, :]$
    $\text{Q}_L = \text{CombQT(NormQuaternion}(\text{Q}_1 \odot \text{Q}_2))$
    $\text{Q}_L = \text{permute}(\text{Q}_L)$ $[Batch, 4, L, L]$
    **return** $\text{Q}_L$
**def** CompDistMap(T $\in [Batch, K, L, 3]$):
    Dist = T[...,None,:] - T[...,None,:,:]
    Dist = $\|\text{Dist}\|$
    Dist = CombQT(Dist)
    DistMap = Sigmoid(Dist - [8, 10, ..., 42])
    **return** DistMap $[Batch, 17, L, L]$
$\text{C}_\alpha\text{DistMap} = \text{CompDistMap(T)}$
$\text{C}_\beta = \text{T} + \text{TranslationRotation}([-0.537, -0.769, -1.208], \text{Q})$
$\text{C}_\beta\text{DistMap} = \text{CompDistMap}(\text{C}_\beta)$
$\text{Translation}_{\text{encode}} = \text{Translation\_K2L(Q, T)}$
$\text{Quaternion}_{\text{encode}} = \text{Quaternion\_K2L(Q)}$
$\text{x2D}' = \text{Cat}(\text{x2D}, \text{C}_\alpha\text{DistMap}, \text{C}_\beta\text{DistMap}, \text{Translation}_{\text{encode}}, \text{Quaternion}_{\text{encode}})$ $[Batch, 196, L, L]$
x2D′ = Conv2d(Norm$\underbrace{\text{(DropOut(LeakyRelu(Conv2d(Norm(x2D}')))))}_{\times 2}$)
$\text{x2D} = \text{x2D} + \text{x2D}'$
**return** x2D

###### 4.2.2 Decoder

Again, the module is divided into two parts: the 1D decoder that utilizes the 2D feature to update the 1D embedding (Algorithm 9) and the structure encoder that predicts new structural information from the 2D feature by taking the existing structural information as queries (Algorithm 10, see Figure S12). In the original structural information, the translation component needs to be multiplied by a series of scale factors such as 1/10, 1/20, ... , 1/100. The scale factor is used to address the issue of excessively large numerical ranges in predicted coordinates, which are not conducive for the network to encode these coordinates effectively. As illustrated in Figure S13, applying different scale factors to the original coordinate values (including $x$, $y$, and $z$ coordinates) allows for the transformation of these values to the linear range of the Sigmoid activation function, thereby facilitating the embedding of coordinate values into 2D feature more conveniently. Notably, the translations predicted from the structure decoder are treated as additional variables and added to the original translations to obtain new translation values, while the quaternions are replaced by the predicted values.

**Algorithm 9** 1DDecoder

**def** 1DDecoder(
  x1D $\in [Batch, M, L, E_1]$,
  x2D $\in [Batch, E_2, L, L]$):

1: AttenMap = Conv2d(Norm$(\underbrace{\text{LeakyRelu(Conv2d(Norm(x2D))))}}_{\times 3}$
2: AttenMap = Softmax(Symmetrization(AttenMap))
3: x1D′ = Linear(LeakyReLU(DropOut(Linear(LayerNorm(x1D)))))
4: x1D = LayerNorm(x1D + x1D′ @ AttenMap)
5: **return** x1D

**Algorithm 10** StructureDecoder

**def** StructureDecoder(
  T $\in [Batch, K, L, 3]$),
  Q $\in [Batch, K, L, 4]$)
  x2D $\in [Batch, E_2, L, L]$
  AnchorList $\in [Batch, K]$ ):

1: $\text{x2D}_{\text{encode}} = \text{x2D}[\text{AnchorList}]$ $\qquad [Batch, K, E_2, L, L]$
2: $\text{T}_{\text{encode}} = \text{Sigmoid}(\text{T} * [1/10, 1/20, ..., 1/100]$
3: $\text{Encode} = \text{x2D}_{\text{encode}} \| \text{T}_{\text{encode}} \| \text{Q}$
4: $\text{T}', \text{Q}' = \text{Conv2d}(\text{Norm}(\text{LeakyReLU}(\text{DropOut}(\text{Conv2d}(\text{Norm}(\text{Encode}))))))$
5: $\text{Q} = \text{QuaternionNorm}(\text{Q}')$
6: $\text{T} = \text{T} + \text{T}'$
7: **return** T, Q

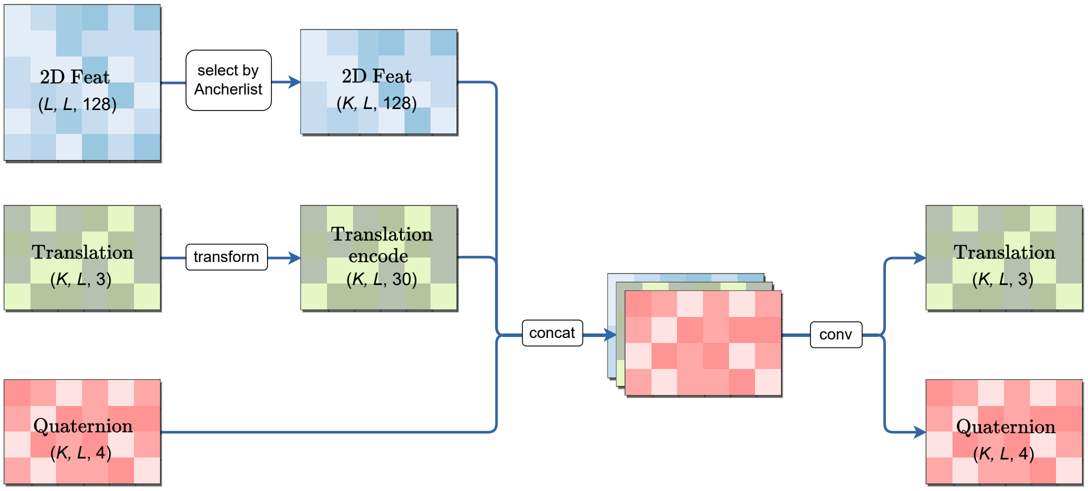


**Figure S12. Schematic architecture of the structure decoder.**

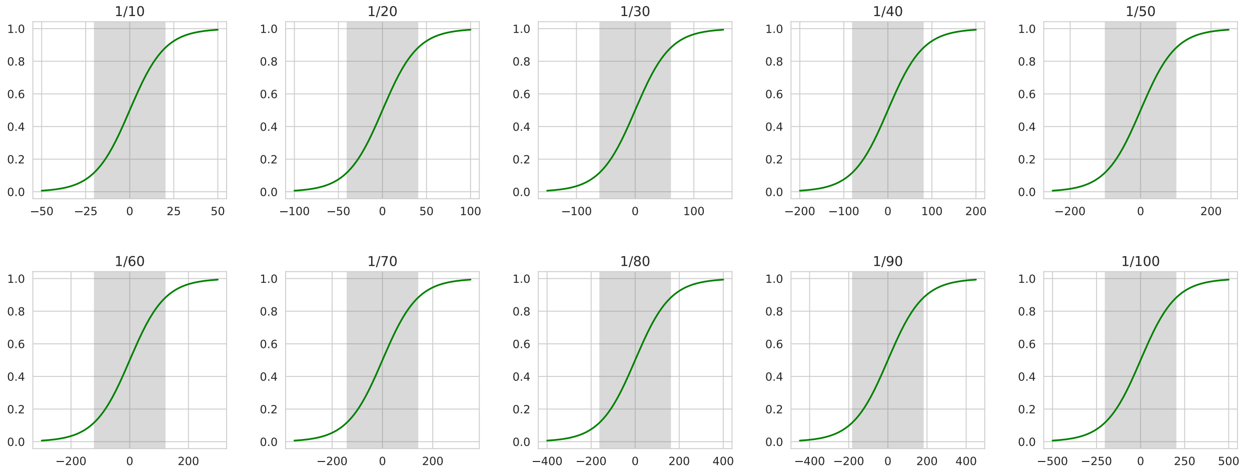


**Figure S13. The scale-factor operation can transform coordinate values into a reasonable range.** The horizontal axis represents the original coordinate values, while the vertical axis displays the values normalized between 0 and -1 after transformation through the scale factor and the Sigmoid activation function. The shaded area denotes the linear range of the Sigmoid activation function, which is more sensitive to values within this region. By combining multiple scale factors, it is possible to encode coordinate values across multiple numerical ranges.

###### 4.2.3 Path Synthesis Attention (PSA) module

The above contents lay the foundation for the transformation between three residues represented by the triplet of $\{i, k, j\}$. Given the known translation and rotation of residue $k$ in the local coordinate system of residue $i$, as well as the translation and rotation of residue $j$ in the local coordinate system of residue $k$, certain relationships can be derived.

To efficiently perform such transformations in a neural network, the SEAT (Select, Extend by AnchorList and Tile) module is designed. The SEAT module consists of two parts (Algorithm 11, see Figure S14). The first part, "Select, Extend by AnchorList," selects the corresponding latent variables based on an AnchorList and duplicates them to obtain the transformation from $i$ to $k$. The second part, "Tile," simply duplicates the variables $K$ times to obtain the transformation from $k$ to $j$.

**Algorithm 11** SelectExtend by AnchorList, and Tile (SEAT)

**def** SEAT(x $\in [Batch, K, L, D]$, AnchorList $\in [K]$):

1: $\mathrm{x}_{i\to k} = \mathrm{x}[:, :, \text{AnchorList}, :]$ Select $\mathrm{x}_{i\to k} \in [Batch, K, K, D]$

2: $\mathrm{x}_{i\to k} = \text{Repeat}(\mathrm{x}_{i\to k}) * L$ Extend $\mathrm{x}_{i\to k} \in [Batch, K, K, L, D]$

3: $\mathrm{x}_{k\to j} = \text{Repeat}(\mathrm{x}) * K$ Tile $\mathrm{x}_{k\to j} \in [Batch, K, K, L, D]$

4: **return** $\mathrm{x}_{i\to k}, \mathrm{x}_{k\to j}$

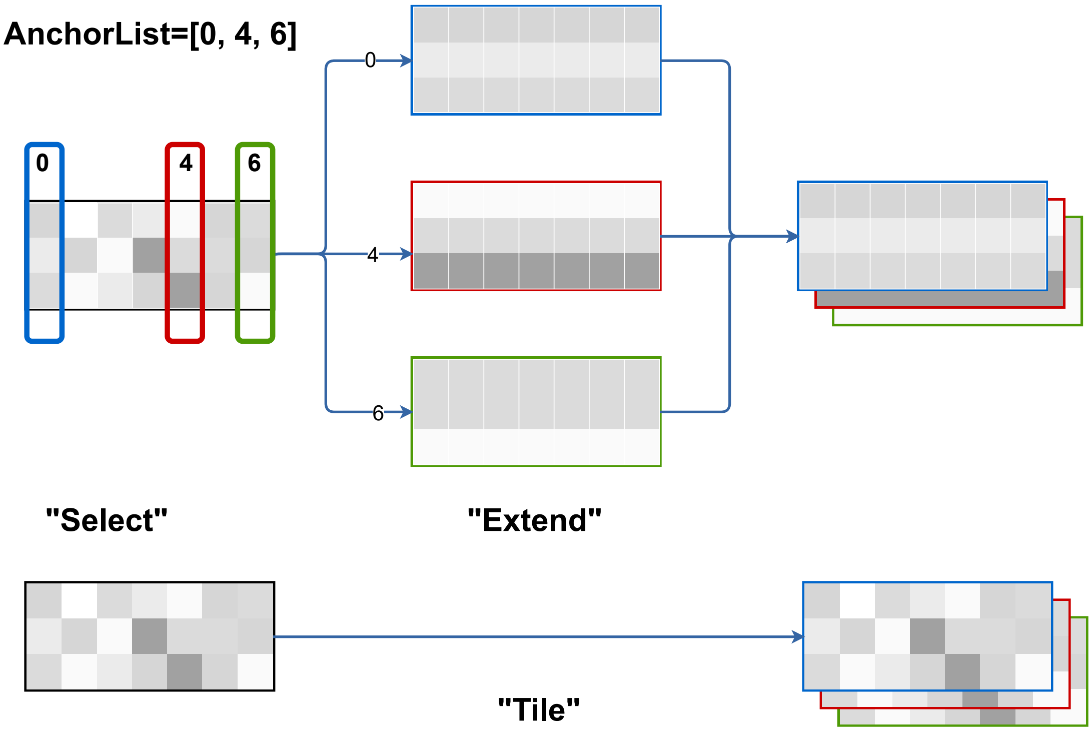


**Figure S14. Schematic presentation of SEAT.**

The detailed implementation of the PSA module is shown in Algorithm 12 and Figure S15. During the path assembly process, for the rotation path of $i \to k \to j$, 2D feature items of each triplet $\{i, k, j\}$ are combined with the corresponding quaternion items to compose a vector representation of path. The weight of each rotation path is estimated by the attention mechanism over the vector representations of all paths (see Figure S16 for a schematic description), and all quaternions are then merged based on their weights. A similar weight calculation method is applied to calculate the weight of each translation path, and the translations are then merged accordingly. Notably, the merging of translation paths requires the use of quaternion $\mathrm{Q}_{ik}$ to rotate the $k \to j$ branch of the translation to the local coordinate system of residue $i$.

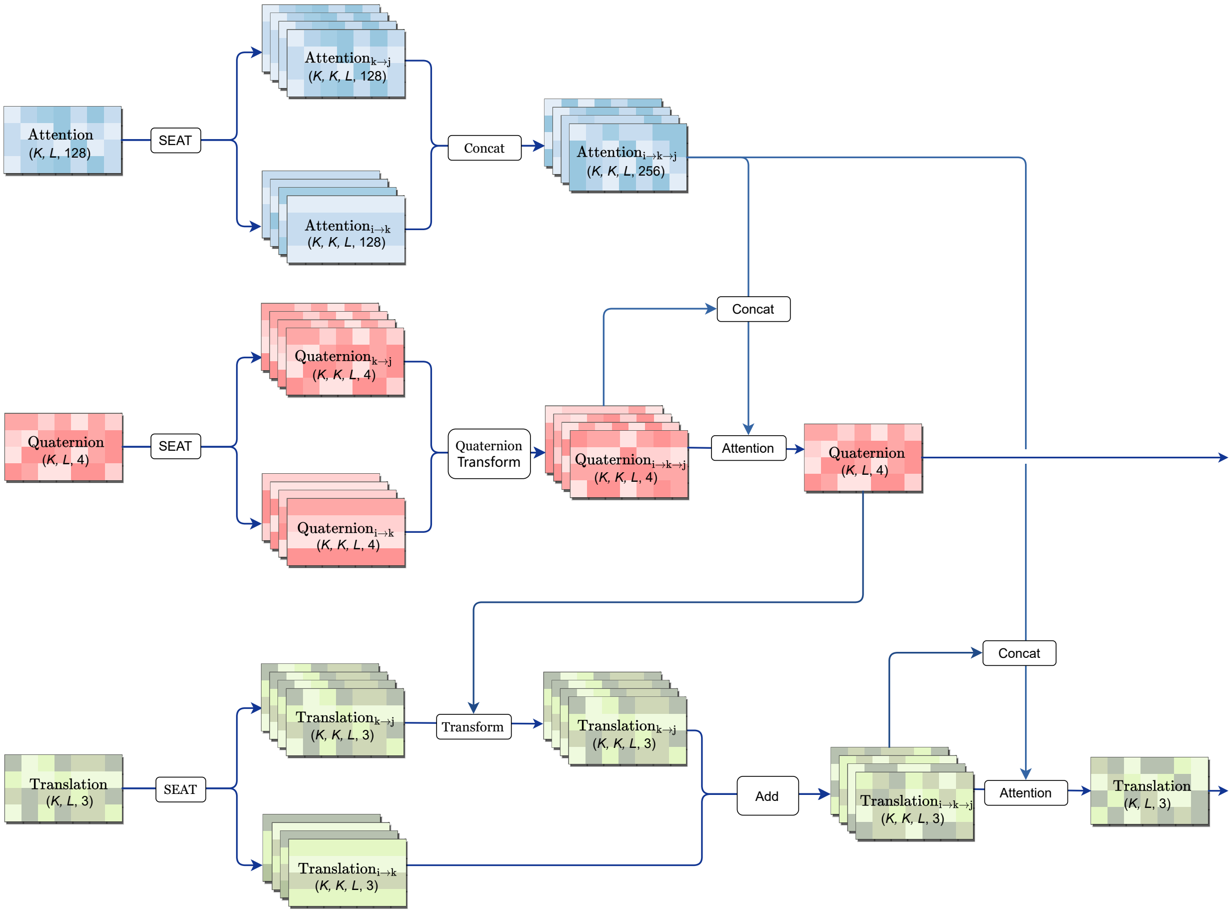


**Figure S15. Schematic architecture of the PSA module.**

**Algorithm 12** PSA

**def** PSA(

AnchorList $\in [Batch, K]$

$\mathrm{T} \in [Batch, K, L, 3]$),

$\mathrm{Q} \in [Batch, K, L, 4]$)

x2D $\in [Batch, E2, L, L]$ ):

**def** AttenWeight(x $\in$ [..., $K$, ...]):

$\mathrm{x} = \mathrm{LayerNorm(x)}$

$\mathrm{q, k} = \mathrm{Linear(x), Linear(x)}$

$\mathrm{Atten} = \mathrm{q} * \mathrm{k/sqrt(q.dim}[-1])$

$\mathrm{Atten} = \mathrm{Softmax(Atten)}$

**return** Atten

Path Synthesis Attention Embedding

$\mathrm{Atten} = \mathrm{x2D[AnchorList]}$ $[Batch, K, L, E_2]$

$\mathrm{Atten}_{i\to k}, \mathrm{Atten}_{k\to j} = \mathrm{SEAT(Atten)}$ $\mathrm{Atten}_{i\to k}, \mathrm{Atten}_{k\to j} \in [Batch, K, K, L, E_2]$

$\mathrm{Atten}_{i\to k\to j} = \mathrm{Atten}_{i\to k} \| \mathrm{Atten}_{k\to j}$ $\mathrm{Atten}_{i\to k\to j} \in [Batch, K, K, L, 2 * E_2]$

Quaternion Synthesis

$\mathrm{Q}_{i\to k}, \mathrm{Q}_{k\to j} = \mathrm{SEAT(Q)}$ $\mathrm{Q}_{i\to k}, \mathrm{Q}_{k\to j} \in [Batch, K, K, L, 4]$

$\mathrm{Q}_{i\to k\to j} = \mathrm{NormQuaternion}(\mathrm{Q}_{i\to k} \odot \mathrm{Q}_{k\to j})$ $\mathrm{Q}_{i\to k\to j} \in [Batch, K, K, L, 4]$

$\omega_{\mathrm{Q}} = \mathrm{Q}_{i\to k\to j} \| \mathrm{Atten}_{i\to k\to j}$ $\omega_{\mathrm{Q}} \in [Batch, K, K, L, 4 + 2 * E_2]$

$\omega_{\mathrm{Q}} = \mathrm{AttenWeight}(\omega_{\mathrm{Q}})$ $\omega_{\mathrm{Q}} \in$ [..., $K$, $K$, ...]

$\mathrm{Q} = \mathrm{NormQuaternion}(\sum(\mathrm{LieAlgebraInterpolation}(\mathrm{Q}_{i\to k\to j}, \omega_{\mathrm{Q}})))$

$\mathrm{Q\_consistence} = \|\mathrm{Q}_{i\to k\to j} - \mathrm{Q.detach}\|^2_{\mathrm{L}_2}$

Translation Synthesis

$\mathrm{T}_{i\to k}, \mathrm{T}_{k\to j} = \mathrm{SEAT(T)}$ $\mathrm{T}_{i\to k}, \mathrm{T}_{k\to j} \in [Batch, K, K, L, 3]$

$\mathrm{T}_{k\to j} = \mathrm{TranslationRotation}(\mathrm{T}_{k\to j}, \mathrm{Q})$

$\mathrm{T}_{i\to k\to j} = \mathrm{T}_{i\to k} + \mathrm{T}_{k\to j}$ $\mathrm{T}_{i\to k\to j} \in [Batch, K, K, L, 3]$

$\omega_{\mathrm{T}} = \mathrm{Sigmoid}(\mathrm{T}_{i\to k\to j}) \| \mathrm{Atten}_{i\to k\to j}$ $\omega_{\mathrm{T}} \in [Batch, K, K, L, 3 + 2 * E_2]$

$\omega_{\mathrm{T}} = \mathrm{AttenWeight}(\omega_{\mathrm{T}})$ $\omega_{\mathrm{T}} \in$ [..., $K$, $K$, ...]

$\mathrm{T} = \sum(\mathrm{T}_{i\to k\to j} \times \omega_{\mathrm{T}})$

$\mathrm{T\_consistence} = \|\mathrm{T}_{i\to k\to j} - \mathrm{T.detach}\|_{\mathrm{L}_2}$

**return** T, Q, T_consistence, Q_consistence

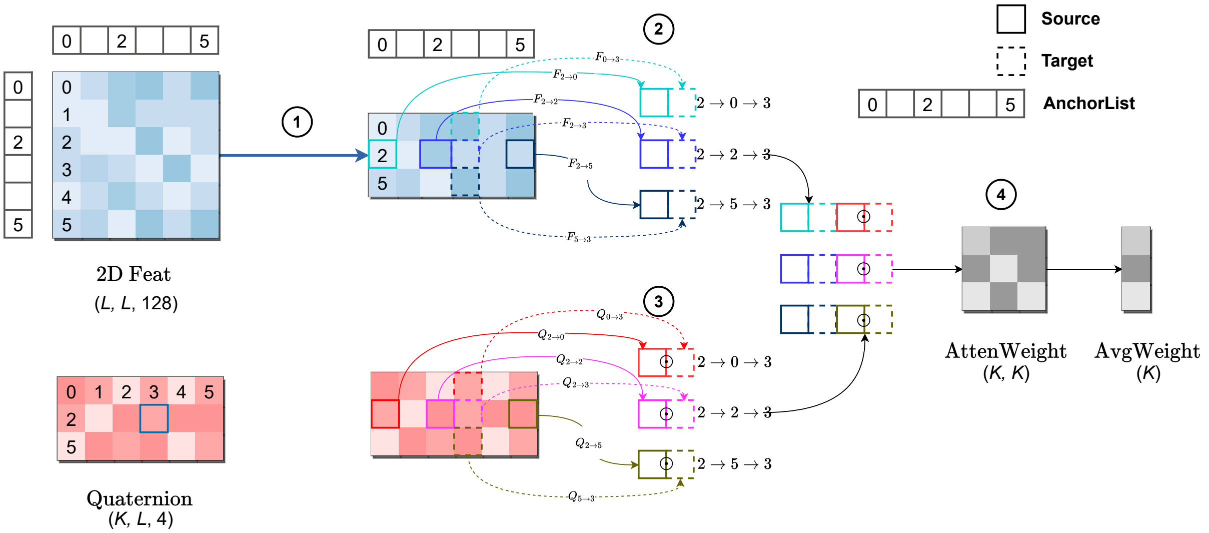


**Figure S16. Schematic description on the stepwise calculation of the PSA attention weights.** In this example, the anchor list contains residues 0, 2 and 5. Suppose that we focus on the rotation of residue 3 (target) relative to the local coordinate system of residue 2 (source). Hence, there exists 3 rotation paths, including $2 \to 0 \to 3$, $2 \to 2 \to 3$ and $2 \to 5 \to 3$. At the first step, the rows corresponding to the anchor residues are collected from the tensor of 2D feature. At the second step, items in the collected features are further recombined to derive the feature representations for all available paths. For instance, 2D feature item [2, 0] (for row 2 and column 0, denoted as $F_{2\to 0}$) and item [0, 3] (for row 0 and column 3, denoted as $F_{0\to 3}$) will be used for predicting the translation and quaternion of residue 0 relative to the frame of residue 2 as well as those of residue 3 relative to the frame of residue 0, respectively. Hence, for the path of $2 \to 0 \to 3$, these items are concatenated as a feature representation of the path. At the third step, the corresponding items are recombined in a similar way (but following the quaternion composition) to derive the quaternion representations of all available paths. At the fourth step, the feature representations and quaternion representations are concatenated, which are used to derive the attention weights between all pairs of paths through the conventional attention mechanism. After averaging, the final weights quantify the importance of each individual rotation path. Attention weights for the translation paths are derived in a similar approach, except that the translation representations (following the translation-rotation coupled operation) are engaged instead of the quaternion representations.

#### 4.3 Other outputs

Cerebra incorporates additional model outputs to stabilize the model training process. The module predicting mask token (Algorithm 13) and the module predicting the distance between $C_\beta$ atoms (Algorithm 14) are connected after the Evoformer Module. The module predicting the plDDT (Algorithm 15) is connected after the Structure Module, along with the module predicting the main-chain dihedral angles (Algorithm 16).

**Algorithm 13** MLM

**def** MLM(x1D $\in [Batch, M, L, E_1]$ from Evoformer):

1: **return** $\text{Linear}(\text{x1D})$ $[Batch, M, L, 23]$

**Algorithm 14** CB dist

**def** PredDist(x2D $\in [Batch, E_2, L, L]$ from Evoformer):

1: Dist = $\text{Conv2d}(\text{LeakyReLU}(\text{Conv2d}(\text{Conv2d}(\text{x2D}))))$ $[Batch, 36, L, L]$

2: **return** $\text{permute}(\text{Dist})$ $[Batch, L, L, 36]$

**Algorithm 15** plDDT

**def** plDDTModule(

x2D $\in [Batch, E_2, L, L]$ from Structure,

T $\in [Batch, K, L, 3]$ from Structure):

1: $\text{Pair} = \text{Permute}(\text{x2D}) + \text{Transpose}(\text{Permute}(\text{x2D}), 1, 2)$ $[Batch, L, L, E_2]$

2: $\text{Dist} = \|\text{T}[..., \text{None}, :] - \text{T}[..., \text{None}, :, :]\|_{L_2}$ $[Batch, K, L, L]$

3: $\text{Cutoff} = [1, 1.5, \ldots, 32.5]$ $[64]$

4: $\text{DistBin} = \text{Cast}(\text{Dist}[..., \text{None}] \leq \text{Cutoff})$ $[Batch, K, L, L, 64]$

5: $\text{Pair} = \text{Repeat}(\text{Pair})$ $[Batch, K, L, L, E_2]$

6: $\text{Pair} = \text{LayerNorm}(\text{Cat}(\text{Pair}, \text{DistBin}))$

7: $\text{Pair} = \text{RELU}(\text{Linear}(\text{ReLU}(\text{Linear}(\text{Pair}))))$ $[Batch, K, L, L, 128]$

8: $\text{Logits} = \text{Linear}(\text{Pair})$ $[Batch, K, L, L, 50]$

9: $\text{Prob} = \text{Softmax}(\text{Logits}, \text{dim} = -1)$

10: $\text{BinCenter} = (\text{arange}(50) + 0.5)/50$ $[50]$

11: $\text{plDDT} = \text{Sum}(\text{Prob} \times \text{BinCenter})$ $[Batch, K, L, L]$

12: **return** $\text{Mean}(\text{plDDT})$ $[Batch, K, L]$

**Algorithm 16** Dihedral Angle

**def** DihedralAngle(x1D $\in [Batch, M, L, E_1]$ from Structure):

1: $\text{Target} = \text{x1D}[:, 0]$ $[Batch, L, E_1]$

2: $\text{Target} = \text{Permute\&Reshape}(\text{Target})$ $[Batch, 1, E_1, L]$

3: **return** $\text{Conv2d}(\text{LeakyReLU}(\text{Conv2d}(\text{Target})))$ $[Batch, 2, 1, L - 1]$

These additional modules help improve the stability and accuracy of the model performance during training. The former two modules assist in predicting the relative positions of the mask tokens and the $C_\beta$ distance matrix, while the latter two modules focus on estimating the local quality of the protein structures predicted. The inclusion of these modules enhances the overall performance and reliability of the Cerebra model.

### 5 Loss Function

#### 5.1 Combination of loss functions

At different stages of training, we adopt different combinations of loss functions to achieve fast convergence of the model.

In the initial training stage:

$$\mathcal{L}_{\text{init}} = 0.3\mathcal{L}_{\text{dist}} + 2\mathcal{L}_{\text{MSA}} + 2\mathcal{L}_{\text{quaternion\_simple}} + 2\mathcal{L}_{\text{translation\_simple}} + 0.1\mathcal{L}_{\text{dihedral}} + 0.01\mathcal{L}_{\text{lDDT}}. \tag{E13}$$

In the large-scale training stage:

$$\begin{aligned}\mathcal{L} =& 0.3\mathcal{L}_{\text{dist}} + 2\mathcal{L}_{\text{MSA}} + 2\mathcal{L}_{\text{quaternion}} + 2\mathcal{L}_{\text{translation}} + 0.1\mathcal{L}_{\text{dihedral}} + 0.01\mathcal{L}_{\text{lDDT}} \\ &+ 0.01\mathcal{L}_{\text{plDDT}} + 0.01\mathcal{L}_{\text{Q\_consistence}} + 0.01\mathcal{L}_{\text{T\_consistence}}\end{aligned}. \tag{E14}$$

In the model fine-tuning stage:

$$\begin{aligned}\mathcal{L}_{\text{fine-tuning}} =& 0.3\mathcal{L}_{\text{dist}} + 2\mathcal{L}_{\text{MSA}} + 2\mathcal{L}_{\text{quaternion}} + 2\mathcal{L}_{\text{translation}} + 0.1\mathcal{L}_{\text{dihedral}} + 0.01\mathcal{L}_{\text{lDDT}} \\ &+ 0.01\mathcal{L}_{\text{plDDT}} + 0.01\mathcal{L}_{\text{Q\_consistence}} + 0.01\mathcal{L}_{\text{T\_consistence}} \\ &+ 0.2\mathcal{L}_{\text{FAPE}} + 0.01\mathcal{L}_{\text{angle}} + 0.01\mathcal{L}_{\text{viol}}\end{aligned}. \tag{E15}$$

The above loss functions can be roughly divided into two parts: the $\mathcal{L}_{\text{dist}}$ and $\mathcal{L}_{\text{MSA}}$ functions used for the Evoformer Module, and the remaining loss functions used for structural optimization.

#### 5.2 Evoformer loss function

$\mathcal{L}_{\text{dist}}$: We use the cross-entropy to evaluate the difference between the predicted distance distribution of $C_\beta$ atoms and the true distance distribution. We divide the distance between $C_\beta$ atoms in the target structure into 36 bins: distances of < 4 Å or > 21Å correspond to 2 bins, respectively, whereas distances between 4 Å and 21 Å are discretized into 34 bins with a bin size of 0.5 Å.

$$\mathcal{L}_{\text{dist}} = \text{mean}(-\sum_{b=1}^{36} y^b \log p^b) \tag{E16}$$

$\mathcal{L}_{\text{MSA}}$: Similar to AlphaFold2, we apply a 15% random mask to the MSA and use a neural network to predict the masked residues:

$$\mathcal{L}_{\text{MSA}} = \text{mean}(-\sum_{c=1}^{22} y^c \log p^c). \tag{E17}$$

In Formula E17, $y$ and $p$ represent the true labels and predicted logits, respectively. For the masked residues, their ground truths can be divided into 22 categories, which include the 20 types of common amino acids, the uncommon amino acids, and the gap token.

#### 5.3 Structure loss function

**Translation Loss**: In Cerebra, both translation and rotation are computed for each residue, where the translation itself represents the $C_\alpha$ coordinates of the residue. We can calculate the coordinates of C, N, and $C_\beta$ atoms using statistical values and quaternions. In the initial training stage, we minimize the difference between the predicted translation and the ground truth coordinates to ensure that each residue is correctly positioned (Algorithm 17).

**Algorithm 17** Comp Translation Simple Loss

**def** $\mathcal{L}_{\text{translation\_simple}}(\tilde{\mathrm{T}} \in [K, L, 3], \tilde{\mathrm{Q}} \in [K, L, 4], \mathrm{Q} \in [L, 4], \mathrm{T}[\mathrm{C}_\alpha, \mathrm{C}, \mathrm{N}, \mathrm{C}_\beta] \in [L, 4, 3])$:

$$\tilde{\mathrm{T}}^{\text{Atom}} = [\tilde{\mathrm{C}}_\alpha, \tilde{\mathrm{C}}, \tilde{\mathrm{N}}, \tilde{\mathrm{C}}_\beta] = \tilde{\mathrm{T}} + \begin{pmatrix} 0. & 0. & 0. \\ 1.523 & 0. & 0. \\ -0.518 & 1.364 & 0. \\ -0.537 & -0.769 & -1.208 \end{pmatrix} \odot \tilde{\mathrm{Q}}$$

$\mathcal{L}_{\text{translation}} = \dfrac{1}{K} \sum_{n=1}^{K} (\min((\|\tilde{\mathrm{T}}^{\text{Atom}} - \mathrm{T}\|, 10)))/L$

**return** $\mathcal{L}_{\text{translation}}$

In both the large-scale training stage and the finetuning stage, we use the multi-structure FAPE loss (Algorithm 18) to guide the residues to move to the correct positions. Similar to the FAPE loss in AlphaFold2[3], we transform the $K$ sets of coordinates to a local coordinate system and calculate their deviations from the coordinates of the corresponding residues in the real protein structure transformed to the same local coordinate system. We also employ the AlphaFold2 method by setting a clipping threshold of 10 Å. That is, if the predicted coordinates differ from the ground truth coordinates by more than 10 Å, we use 10 Å as the deviation.

**Algorithm 18** Comp Translation Loss

**def** $\mathcal{L}_{\text{translation}}(\tilde{\mathrm{T}} \in [K, L, 3], \tilde{\mathrm{Q}} \in [K, L, 4], \mathrm{Q} \in [L, 4], \mathrm{T}[\mathrm{C}_\alpha, \mathrm{C}, \mathrm{N}, \mathrm{C}_\beta] \in [L, 4, 3])$:

$\mathrm{T}_{ij} = \mathrm{T}_i \odot \mathrm{Q}_j^{-1}$

$\mathrm{T}_{ij} = \mathrm{T}_{ij} - \mathrm{T}_{ii}$

$\tilde{\mathrm{Q}}_{ij|k} = \tilde{\mathrm{Q}}_{ki} \odot \bar{\mathrm{Q}}_{kj}^{-1}$

$$\tilde{\mathrm{T}}_{ij|k}^{\text{Atom}} = \begin{pmatrix} 0. & 0. & 0. \\ 1.523 & 0. & 0. \\ -0.518 & 1.364 & 0. \\ -0.537 & -0.769 & -1.208 \end{pmatrix} \odot \tilde{\mathrm{Q}}_{ij|k}$$

$\tilde{\mathrm{T}}_{ij|k} = \tilde{\mathrm{T}}_{ki} \odot \bar{\mathrm{Q}}_{kj}^{-1}$

$\tilde{\mathrm{T}}_{ij|k} = \tilde{\mathrm{T}}_{ij|k} - \tilde{\mathrm{T}}_{ii|k}$

$\tilde{\mathrm{T}}_{ij|k} = [\tilde{\mathrm{C}}_{\alpha ij|k}, \tilde{\mathrm{C}}_{ij|k}, \tilde{\mathrm{N}}_{ij|k}, \tilde{\mathrm{C}}_{\beta_{ij|k}}] = \tilde{\mathrm{T}}_{ij|k} + \tilde{\mathrm{T}}_{ij|k}^{\text{Atom}}$

$\mathcal{L}_{\text{translation}} = \dfrac{1}{K} \sum_{k=1}^{K} (\min((\|\tilde{\mathrm{T}}_{ij|k} - \mathrm{T}_{ij}\|, 10)))/L^2$

**return** $\mathcal{L}_{\text{translation}}$

**Quaternion Loss**: In the initial training stage, we only calculate the quaternions of residues with respect to anchor residues and use the $L_1$ loss to compute the difference between the predicted values and the ground truth (Algorithm 19).

**Algorithm 19** Comp Quaternion Simple Loss

**def** $\mathcal{L}_{\text{quaternion\_simple}}(\tilde{\mathrm{Q}} \in [K, L, 4], \mathrm{Q} \in [L, 4])$:

$\mathcal{L}_{\text{quaternion}} = \frac{1}{K} \sum_{k=1}^{K} (\text{abs}(\tilde{\mathrm{Q}} - \mathrm{Q}))/L$

**return** $\mathcal{L}_{\text{quaternion}}$

In the large-scale training stage and finetuning stage, we use Algorithm 20 to compute the difference between the predicted quaternions and the ground truth along different rotation paths.

**Algorithm 20** Comp Quaternion Loss

**def** $\mathcal{L}_{\text{quaternion}}(\tilde{\text{Q}} \in [K, L, 4], \text{Q} \in [L, 4])$:

1: $\text{Q}_{ij} = \text{Q}_i \odot \text{Q}_j^{-1}$
2: $\tilde{\text{Q}}_{ij|k} = \tilde{\text{Q}}_{ki} \odot \bar{\text{Q}}_{kj}^{-1}$
3: $\mathcal{L}_{\text{quaternion}} = \frac{1}{K}\sum_{k=1}^{K}(\text{abs}(\tilde{\text{Q}}_{ij|k} - \text{Q}_{ij}))/L^2$
4: **return** $\mathcal{L}_{\text{quaternion}}$

We use the $L_2$ norm to calculate the variation of translation updated along different paths (Formula E18), enforcing each path to be closer to the desired result.

$$\mathcal{L}_{\text{T\_consistence}} = \text{mean}(\text{T\_consistence}) \tag{E18}$$

Similarly, we calculate the path divergence loss function for quaternions (Formula E19). This allows us to measure the deviation of the rotation paths and guide them towards the desired outcome.

$$\mathcal{L}_{\text{Q\_consistence}} = \text{mean}(\text{Q\_consistence}) \tag{E19}$$

To evaluate the prediction model, we adopt the metric called predicted local distance difference test (plDDT), which estimates the local quality of a protein structure prediction. During the training process, we use the true local distance difference test (lDDT) as the supervision signal and train the plDDT using the $L_1$ loss (Formula E20).

$$\mathcal{L}_{\text{plDDT}} = \text{mean}(\text{abs}(\text{plDDT} - \text{lDDT})) \tag{E20}$$

Cerebra calculates the main-chain dihedral angles $\phi$ and $\psi$ between consecutive residues to constrain the relative torsion between adjacent residues. The label values and the method to compute the loss for $\phi$ and $\psi$ can be found in Algorithm 21.

In the finetuning stage of the model, we introduce a side-chain prediction module as well as $\mathcal{L}_{\text{FAPE}}$, $\mathcal{L}_{\text{angle}}$ and the structural clash loss term $\mathcal{L}_{\text{viol}}$ from AlphaFold2 into the training process. The $\mathcal{L}_{\text{angle}}$ and $\mathcal{L}_{\text{viol}}$ are set to be consistent with AlphaFold2, while $\mathcal{L}_{\text{FAPE}}$ undergoes minor adjustments. For each structure, $\mathcal{L}_{\text{FAPE}}$ is computed, and then the average is taken.

**Algorithm 21** $\mathcal{L}_{\text{dihedral}}$

**def** CompDihedral($\tilde{\psi}, \tilde{\phi}$, $\mathrm{C}_\alpha, \mathrm{C}, \mathrm{N} \in [L, 3]$):

$\mathrm{a} = \mathrm{N}[1:] - \mathrm{C}[:-1]$
$\mathrm{b} = \mathrm{C} - \mathrm{C}_\alpha$
$\mathrm{c} = \mathrm{N} - \mathrm{C}_\alpha$
$\mathrm{ab} = \mathrm{a} \times \mathrm{b}[:-1]$
$\mathrm{bc} = \mathrm{b}[:-1] \times \mathrm{c}[:-1]$
$\mathrm{ca} = \mathrm{c}[:-1] \times \mathrm{a})$
$\mathrm{cos\_ca\_b} = \sum(\mathrm{ca} * \mathrm{b}[:-1])/(\|\mathrm{ca}\| * \|\mathrm{b}[:-1]\|)$
$\mathrm{cospsi} = \pi - \arccos(\sum(\mathrm{ab} * \mathrm{bc})/(\|\mathrm{ab}\| * \|\mathrm{bc}\|))$
$\psi = \mathrm{sign}(\mathrm{cos\_ca\_b}) * \mathrm{cospsi}$
$\mathrm{d} = \mathrm{C}[:-1] - \mathrm{N}[1:]$
$\mathrm{bc} = \mathrm{b}[1:] \times \mathrm{c}[1:]$
$\mathrm{cd} = \mathrm{c}[1:] \times \mathrm{d}$
$\mathrm{bd} = \mathrm{b}[1:] \times \mathrm{d}$
$\mathrm{cos\_bd\_c} = \sum(\mathrm{bd} * \mathrm{c}[1:])/(\|\mathrm{bd}\| * \|\mathrm{c}[1:]\|)$
$\mathrm{cosphi} = \pi - \arccos(\sum(\mathrm{bc} * \mathrm{cd})/(\|\mathrm{bc}\| * \|\mathrm{cd}\|))$
$\phi = \mathrm{sign}(\mathrm{cos\_bd\_c}) * \mathrm{cosphi}$
$\mathcal{L}_{\text{dihedral}} = \frac{1}{L-1} \sum(\mathrm{abs}(\tilde{\psi} - \psi)) + \frac{1}{L-1} \sum(\mathrm{abs}(\tilde{\phi} - \phi))$

### 6 Training of Cerebra and OpenFold

#### 6.1 Dataset and training strategy

We initially trained our model using the OpenFold dataset, which consisted of approximately 116,000 proteins in total after excluding small proteins of length < 50 residues (see **Methods**). Notably, the exclusion of protein monomers with the sequence length < 50 residues allowed the model to focus on proteins with a certain level of complexity and structural diversity during training, which is important for accurately predicting protein topology. During different stages of training, we used datasets of varying sizes. Here is a concrete description.

- **10K Dataset**: Randomly select 10,000 monomers with the sequence length between 100 and 400 residues and resolution < 3Å.
- **3Å Dataset**: Use all PDB samples with resolution < 3 Å and length > 50 residues. This dataset contains ~96,000 protein monomers.
- **Entire Dataset**: Use all PDB samples with length > 50 residues. This dataset contains ~116,000 protein monomers.
- **Expanded Dataset**: Besides the PDB structures, this dataset also contains ~268,000 proteins from the AlphaFold2 distillation dataset. Moreover, ~46,000 disordered proteins with AlphaFold2-predicted structures are used for fine-tuning at the late training stage.

The input features for this model can be classified into two types: the MSA and the embedding of the target sequence generated by the ESM-2 (3B) model, where the MSA data are obtained from the OpenFold dataset. We employ different training sets, features and loss functions in different stages to accelerate model training. The details of the training strategies can be found in Table S5.

**Table S5. Training strategy**

| Stage | Datasets | Loss Function | Feature | Anchor Num | Crop Size | Side Chain Prediction |
|---|---|---|---|---|---|---|
| 1 | 10K Dataset | $\mathcal{L}_{\text{init}}$ | MSA | 32 | 256 | ✗ |
| 2 | 10K Dataset | $\mathcal{L}$ | MSA | 24 | 256 | ✗ |
| 3 | 3Å Dataset | $\mathcal{L}$ | MSA | 24 | 256 | ✗ |
| 4 | 3Å Dataset | $\mathcal{L}$ | MSA + ESM-2 | 24 | 256 | ✓ |
| 5 | Entire Dataset | $\mathcal{L}_{\text{fine-tuning}}$ | MSA + ESM-2 | 24 | 384 | ✓ |
| 6 | Expanded Dataset | $\mathcal{L}_{\text{fine-tuning}}$ | MSA + ESM-2 | 24 | 384 | ✓ |

In the training process, the number of anchor residues is selected based on the sequence length. When the length is less than 128, 12 anchor residues are used. When the sequence length is between 128 and 256, 24 anchor residues are used. Here, we denote the sequence length as $L$ and the number of anchor residues as $K$. Therefore, the selection of anchor residues during the training process can be determined by Formula E21:

$$\text{AnchorList} = \text{int}([L/(K+1)] * [1, 2, ..., K] + \text{randint}(0, L/K)). \quad \text{(E21)}$$

#### 6.2 Training process for the ablation experiments

1. All ablation experiments were conducted on the 10K Dataset.

2. For the Cerebra series models, if not specifically mentioned, 32 anchors were used.

3. The learning rate for all model training was set to $10^{-3}$ with a 1000-step warm-up.

4. For OpenFold, the batch size was set as 6. Cerebra, however, used a dynamic capacity batch size. Sequences in the training set were shuffled in advance and grouped into small batches with similar lengths. The shortest sequence length within each batch was used as the cropping length for all sequences. For sequences longer than 256, they were cropped to 256. The batches were composed as follows: [96: 8, 128: 6, 144: 6, 192: 4, 256: 3], where the first number represents the maximum sequence length tolerance and the second number represents the number of sequences in the batch.

5. We used $\mathcal{L}_{\text{init}}$ loss to train our models.

6. All models were trained using PyTorch and in the mixed precision mode.

In the training process of OpenFold, no templates were used and the additional MSA count was 1024.

#### 7 Inference of OpenFold and Cerebra

As Cerebra did not use protein template information during the training process, for the purpose of fair comparison, three sets of OpenFold model parameter weights (without templates), namely finetuning_no_templ_1, finetuning_no_templ_2 and finetuning_no_templ_ptm_1 released for OpenFold, were selected for the inference of OpenFold on the CAMEO datasets. Based on their mean TM-scores on the CAMEO 2024 validation set, we selected five checkpoints as the final model parameters for Cerebra. Each checkpoint predicts $K$ structures, where $K$ is the number of anchors. For each checkpoint, all $K$ structures are superimposed onto the $\mathrm{int}(K/2)^{\mathrm{th}}$ structure and averaged to obtain a checkpoint-level prediction. During the prediction process, Formula E22 is used to select anchor residues, where $L$ denotes the sequence length:

$$\begin{cases} \mathrm{AnchorList} = \mathrm{int}(((L-8)/K) * [1, 2, ..., K]) \\ \mathrm{AnchorList} = \mathrm{clip}(\mathrm{AnchorList}, \mathrm{min} = 2, \mathrm{max} = L - 2) \end{cases} \tag{E22}$$

**Integration of predicted results**: The final prediction is selected from the five checkpoint-level predictions based on the highest mean pLDDT.

### 8 Hallucination-based *de novo* protein design

#### 8.1 MCMC optimization and state retention

Each trajectory was initialized with an amino acid sequence sampled from background residue frequencies derived from BLOSUM62 composition statistics. At MCMC step $t$, a trial sequence was generated by mutating one or more positions in the retained sequence. Cerebra was evaluated on the trial sequence, and the change in the confidence-based loss defined in Equation 5 was calculated as

$$\Delta\mathcal{L}_t = \mathcal{L}(s_{\text{trial}}) - \mathcal{L}(s_{\text{current}}). \tag{E23}$$

The trial sequence was accepted with probability

$$P_{\text{accept}} = \begin{cases} 1, & \Delta\mathcal{L}_t < 0, \\ \exp\left(-\Delta\mathcal{L}_t / T_t\right), & \Delta\mathcal{L}_t \geq 0, \end{cases} \tag{E24}$$

where the simulated-annealing temperature followed

$$T_t = T_0 \times 0.5^{t/h}. \tag{E25}$$

Here, $T_0 = 0.01$ and $h$ denotes the temperature half-life. After an accepted proposal, the retained sequence, loss, residue-level pLDDT profile, and predicted coordinates were replaced by their trial values; after a rejected proposal, all retained quantities remained unchanged. Thus, the pLDDT reported along an optimization trajectory was $1 - \mathcal{L}(s_{\text{current}})$ and did not include confidence values from rejected trials. Accepted structures were written as PDB files during sampling. The terminal output of each trajectory included the final retained structure and sequence together with the protein length, trajectory identifier, last accepted step, and final mean pLDDT.

#### 8.2 Protocol-specific mutation proposals

The Cerebra-hallucination-refined protocol used 5,000 MCMC steps with a temperature half-life ($h$) of 1,000 steps. Its mutation radius was reduced linearly from three simultaneously resampled positions to one over the trajectory. Positions were sampled uniformly without replacement, and the new residue identities were sampled from the same BLOSUM62-derived background distribution used to initialize the sequence. This small mutation radius supported gradual exploration and refinement following an AlphaFold2-based hallucination strategy[4].

The Cerebra-hallucination-fast protocol used 1,500 MCMC steps with a temperature half-life ($h$) of 500 steps. Its mutation radius was reduced linearly from 15 positions to one. Candidate positions were preferentially sampled from the lowest pLDDT quartile of the retained structure, concentrating proposals in locally uncertain regions. To limit the systematic over-selection of flexible termini, the first and last three residues were downweighted during position sampling. At each selected position, the replacement was drawn uniformly from the 19 amino acids other than the current identity. This confidence-guided proposal scheme enabled larger early updates and fewer Cerebra evaluations[5]. Both protocols otherwise used the same confidence objective and Metropolis acceptance rule described above.

#### 8.3 Trajectory generation and production settings

Both protocols were benchmarked independently at sequence lengths of 50, 100, 150 and 200 residues, with 500 trajectories generated for each protocol–length combination. Fifty complete trajectories from each combination were used to summarize the optimization dynamics in Figure S9. Based on the comparison reported in this figure, the refined protocol was used to produce the 50- and 100-residue backbones, whereas the fast protocol was used for the 150- and 200-residue backbones. For each length, the terminal backbones from the first 100 production trajectories were passed to downstream evaluation.

##### 8.4 Sequence design, refolding and novelty evaluation

Designability was evaluated using the unconditional-refolding workflow of Scaffold-Lab[6]. For each generated backbone, ProteinMPNN[7] produced 10 fixed-backbone sequence designs, each of which was independently refolded with ESMFold[8]. Each refolded structure was aligned to its generated target, after which backbone RMSD and TM-score were calculated. The lowest backbone RMSD among the 10 designs was used as the per-backbone summary, together with the corresponding self-consistency TM-score and ESMFold pLDDT. A generated backbone was classified as designable when at least one designed sequence yielded a refolded structure with a backbone RMSD below 2.0 Å.

Structural novelty was evaluated independently of the refolding analysis by querying each generated backbone against PDB structures with Foldseek[9]. For every query with a valid result, we defined the PDB-TM score as the maximum TM-score among its PDB hits. Accordingly, a lower PDB-TM score indicates lower structural similarity to experimentally deposited protein structures.

#### Supplementary References


1. Pedregosa, F., Varoquaux, G., Gramfort, A., *et al.* Scikit-learn: Machine Learning in Python. **12,** 2825–2830. ISSN: 1532-4435. http://jmlr.org/papers/v12/pedregosa11a.html (2011).
2. Mirdita, M., Schütze, K., Moriwaki, Y., *et al.* ColabFold: making protein folding accessible to all. *Nature Methods* **19,** 679–682. ISSN: 1548-7105. https://doi.org/10.1038/s41592-022-01488-1 (2022).
3. Jumper, J., Evans, R., Pritzel, A., *et al.* Highly accurate protein structure prediction with AlphaFold. *Nature* **596,** 583–589. ISSN: 1476-4687. http://dx.doi.org/10.1038/s41586-021-03819-2 (2021).
4. Wicky, B. I. M., Milles, L. F., Courbet, A., *et al.* Hallucinating symmetric protein assemblies. *Science* **378,** 56–61. https://www.science.org/doi/abs/10.1126/science.add1964 (2022).
5. Zhang, B., Liu, K., Zheng, Z., *et al.* Protein language model supervised motif-scaffolding design with GPDL. *International Journal of Biological Macromolecules* **331,** 148441. https://doi.org/10.1016/j.ijbiomac.2025.148441 (2025).
6. Zheng, Z., Zhang, B., Zhong, B., *et al.* Scaffold-Lab: Critical evaluation and ranking of protein backbone generation methods in a unified framework. *PLOS Computational Biology* **22,** e1014290. https://doi.org/10.1371/journal.pcbi.1014290 (2026).
7. Dauparas, J., Anishchenko, I., Bennett, N., *et al.* Robust deep learning-based protein sequence design using ProteinMPNN. *Science* **378,** 49–56. https://doi.org/10.1126/science.add2187 (2022).
8. Lin, Z., Akin, H., Rao, R., *et al.* Evolutionary-scale prediction of atomic-level protein structure with a language model. *Science* **379,** 1123–1130. https://www.science.org/doi/abs/10.1126/science.ade2574 (2023).
9. Van Kempen, M., Kim, S. S., Tumescheit, C., *et al.* Fast and accurate protein structure search with Foldseek. *Nature Biotechnology* **42,** 243–246. https://doi.org/10.1038/s41587-023-01773-0 (2024).
